# Ku-binding motifs in RAG2, XLF, PAXX and MRI support functional redundancy during V(D)J recombination

**DOI:** 10.1101/2023.12.12.570754

**Authors:** Satish K. Tadi, Armelle Gesnik, Philippe Frit, Florence Iehl, Virginie Ropars, Florent Dingli, Damarys Loew, Patrick Calsou, Isabelle Callebaut, Jean-Baptiste Charbonnier, Jean-Pierre de Villartay

**Author notes:** Corresponding authors, Jean-Pierre de Villartay, DGSI; Institut Imagine, 24 bd du Montparnasse, 75015 Paris, France, Satish K. Tadi.

## Abstract

The interaction of several partners with Ku through Ku-binding motifs (KBMs) in their sequences governs their enrolment in NHEJ repair complexes. Here, we first established more specifically the function of KBMs in V(D)J recombination as the molecular basis of functional redundancy between XLF and the NHEJ proteins MRI and PAXX. Then, given the functional redundancy between RAG2 and XLF, we explored the hypothesis of a KBM-mediated interaction between RAG2 and Ku. Through sequence alignment and biophysical methods, we identified a KBM at the C-terminus of RAG2 (R2CT) that mediates its interaction with Ku both *in vitro* and *in cellulo*. Notably, we showed that R2CT/Ku interaction is independent of the RAG nuclease activity. Finally, we demonstrated that the respective KBMs of RAG2 and XLF support their functional redundancy for V(D)J recombination.

## Introduction

In mammalian cells DNA double-strand breaks (DSBs) are considered the most toxic lesions which are mainly repaired by two pathways: homologous recombination (HR) or non-homologous end joining (NHEJ) ^1, 2^. HR functions to repair DSBs in the S and G2 phases of the cell cycle using the sister chromatid as a template for accurate repair while NHEJ is the predominant pathway that is active in all phases of the cell cycle outside mitosis and is the primary pathway of DSB repair in G1-phase cells ^2^. The core factors that are absolutely required for NHEJ include Ku (Ku70/Ku80 heterodimeric complex) which binds to broken DNA ends; DNA-PKcs, the DNA dependent protein kinase catalytic subunit that forms the DNAPK complex in association with Ku; DNA ligase IV (LIG4), which ligates broken DNA ends; XRCC4 (X4), which is essential for the stability and activity of LIG4 and, Cernunnos/XLF, which bridges DNA ends during NHEJ ^1–4^. X4 and XLF interact through their globular head domains and are able to form long filaments *in vitro*. Given this structure, it was proposed that the X4/XLF filament could form a synapse that would facilitate DNA end tethering for their subsequent ligation by NHEJ ^5–10^. Indeed, recent structural studies identified at least two forms of synaptic complexes in which DNA ends are hold together with a minimal XRCC4/XLF complex unit containing one central XLF flanked by one XRCC4 molecule on each side (for a review see ^11–12^).

V(D)J recombination is the process by which B and T lymphocytes randomly assemble variable (V), diversity (D), and joining (J) gene segments to generate unique antigen receptors that can virtually recognize any type of antigens ^13–15^. This process is initiated when the Recombination-Activating Gene products RAG1 and RAG2 (RAG1/2) introduce DSBs between V, D or J coding gene segments and flanking recombination signal sequences (RSS). RAG1/2 cleavage leads to the formation of a pair of hairpin-sealed coding ends (CEs) and a pair of blunt signal ends (SEs) ^15^. Subsequently, NHEJ is the only known process to recognize and efficiently repair these DNA ends forming a coding joint (CJ) and a reciprocal signal joint (SJ) ^16^. Unexpectedly, although XLF function is critical for the repair of genotoxic-induced DSBs in mouse embryonic fibroblasts (MEFs) and embryonic stem cells, it is dispensable for the NHEJ-mediated repair of RAG DSBs in lymphocytes ^10,17^. One prominent difference between genotoxic and V(D)J induced programmed DSBs is the presence of RAG1/2 in the latter. The RAG1/2 complex is known to remain on the DSB it has initiated and to form the post cleavage complex (PCC)^18^. PCC provides a possible means to tether DNA ends, which would be redundant to the expected function of X4/XLF ^19^. Under this hypothesis, the sole presence of the RAG-PCC would provide a DNA repair synapse complementing the absence of the X4/XLF during V(D)J recombination, while such a backup synapsis would be missing at genotoxic-driven DSBs ^19^. The stability of the RAG-PCC relies on the RAG2 C-terminal region (R2CT), a noncore region that is dispensable for V(D)J recombination, as shown *in vitro* and *in vivo* in the *RAG2^c/c^*mouse model specifically engineered to restrict RAG2 to its core region ^20, 21^. V(D)J recombination is not grossly affected in either *RAG2^c/c^* or *XLF* single mutant conditions, but is fully abrogated in *RAG2^c/c^/XLF* mice, resulting in severe combined immune deficient (SCID) animals devoid of mature B and T lymphocytes, highlighting the functional redundancy between the two factors during the DNA repair phase of V(D)J recombination ^22, 23^. Based on these observations, we proposed a two-synapses model in which the RAG-PCC complex on one hand and X4/XLF on the other hand are critical to maintain genome integrity during V(D)J recombination ^9, 10, 19^.

The functional redundancy between XLF and RAG2 during V(D)J recombination was further extended between XLF and several other factors involved in NHEJ such as ATM ^22^, H2AX ^22,3^, 53BP1 ^24^, paralog of XRCC4 and XLF (PAXX) ^25,26,27,28^ and modulator of retrovirus infection homologue (MRI/CYREN) ^29^. A combined deficiency of XLF with either one of these factors leads to abortive V(D)J recombination in lymphoid cells *in vitro* even though the singular loss of any of these proteins does not lead to notable defects. *In vivo*, these double mutant conditions result most often in embryonic lethality caused by massive neuron apoptosis, which is accompanied by impaired immune cells development^27, 28, 30^.

Several accessory factors in the NHEJ process are recruited to DSBs via conserved protein domains or motifs. Three of these conserved modes of interaction comprise Ku-binding motifs (KBMs), FHA domains, and BRCT domains ^31–33^. A short peptide sequence of 10-15 amino acids necessary for binding to Ku was first identified in Aprataxin Polynucleotide Kinase/Phosphatase-like Factor (APLF) and named A-KBM. Then, A-KBM was found to be present in the other DSB repair factors Werner syndrome protein (WRN) and MRI. Distinct motifs necessary for binding to Ku were later identified in the very C-termini of XLF (X-KBM) and PAXX (P-KBM). Regarding the role of KBMs in DNA-ends synapsis, recent structural studies identify KBMs-mediated interactions of XLF and PAXX with Ku80 and Ku70, respectively as part of the protein networks involved in the assembly of various forms of synaptic complex at DSBs DNA ends ^34,35,36^. The importance of the interactions between Ku and the KBMs of XLF and PAXX for DNA ends synapsis was also demonstrated in functional assays in cells ^32, 37^. Moreover, it was reported recently that binding of the small molecule IP6 to Ku adds a further level of regulation to the X-KBM mediated Ku-XLF interaction ^34^.

Given that KBM-mediated interaction of several Ku partners governs their enrolment in NHEJ repair complexes, we first established more specifically KBMs function in V(D)J recombination as the molecular basis of functional redundancy between XLF and the NHEJ proteins MRI and PAXX. Then, given the functional redundancy between RAG2 and XLF, we explored the hypothesis of a KBM-mediated interaction between R2CT and Ku. Through sequence alignment and biophysical analyses, we identified a KBM at the very C-terminus of RAG2 that mediates its interaction with Ku both *in vitro* and *in cellulo*. Notably, we showed that R2CT/Ku interaction is independent from the DNA breaking activity of the RAG complex. Finally, we demonstrated that the respective KBMs of RAG2 and XLF support their functional redundancy for V(D)J recombination.

## Results

### Combined deletion of KBMs in MRI and XLF failed to restore V(D)J activity

To reconstitute V(D)J recombination in cells, Abelson murine leukemia virus transformed pro-B cell-lines were used (*v-abl* pro-B cells), which, upon treatment with the *v-abl* kinase inhibitor (ABLki-PD180970, SIGMA), leads to G1 cell cycle arrest, the induction/stabilization of RAG1/2, and V(D)J recombination of the endogenous immunoglobulin kappa light chain (*Igk*) locus or any chromosomally integrated retroviral recombination substrates ^38, 39^. Cells were transduced with the MX-RSS-GFP/IRES-hCD4 retroviral recombination substrate (pMX-INV) ^40,41^ that allows for GFP expression upon successful chromosomal inversional RAG-mediated recombination (Fig. 1a). Given the reported functional redundancy between MRI and XLF ^29^, CRISPR/Cas9-generated *MRI^-^*^/-^, *MRI^-/-^/XLF^-/-^* and *MRI^-/-^/PAXX^-/-^ v-abl* pro-B cells (Fig.S1a-S1c) were first used to evaluate the role of MRI and XLF respective KBMs through complementation analyses. Analysis of GFP expression showed that ABLki treatment of *XLF^-/-^* or *MRI^-/-^ v-abl* pro-B cells containing pMX-INV leads to V(D)J recombination at similar levels to those observed in WT *v-abl* pro-B cells (Fig. 1b). Likewise, *MRI^-/-^/PAXX^-/-^ v-abl* pro-B cells showed no detectable V(D)J recombination defect, suggesting that MRI and PAXX are epistatic in function. In contrast and in accordance with previous report ^29^, V(D)J recombination was severely impaired in *MRI^-/-^/XLF^-/-^ v-abl* pro-B cells (Fig. 1b), highlighting their functional redundancy. V(D)J recombination was then assayed in *MRI^-/-^/XLF^-/-^*cells upon complementation with XLF^WT^, MRI^WT^, XLF^ΔC^, and MRI^ΔNΔC^ variants, the two latter lacking their known respective KBMs (Fig. 1c). As expected, given their known functional redundancy, complementation with either XLF^WT^ or MRI^WT^ restored V(D)J recombination in *MRI^-/-^/XLF^-/-^ v-abl* pro-B cells, as indicated by GFP expression after ABLki treatment (Fig. 1d). This restoration was dependent upon the presence of their respective KBMs as shown by the decreased complementation when using XLF^ΔC^ or MRI^ΔNΔC^. Furthermore, the co-expression of the two factors lacking their respective KBMs (MRI^ΔNΔC^+ XLF^ΔC^) also failed to fully complement the V(D)J activity, thus emphasizing the role of KBMs (Fig. 1d). These results show that, in G1 phase lymphocytes, the KBMs of MRI and XLF are functionally redundant, arguing that the redundancy between MRI and XLF is likely controlled by their interaction with Ku for the repair of RAG induced DSBs during V(D)J recombination.

**Fig. 1.**
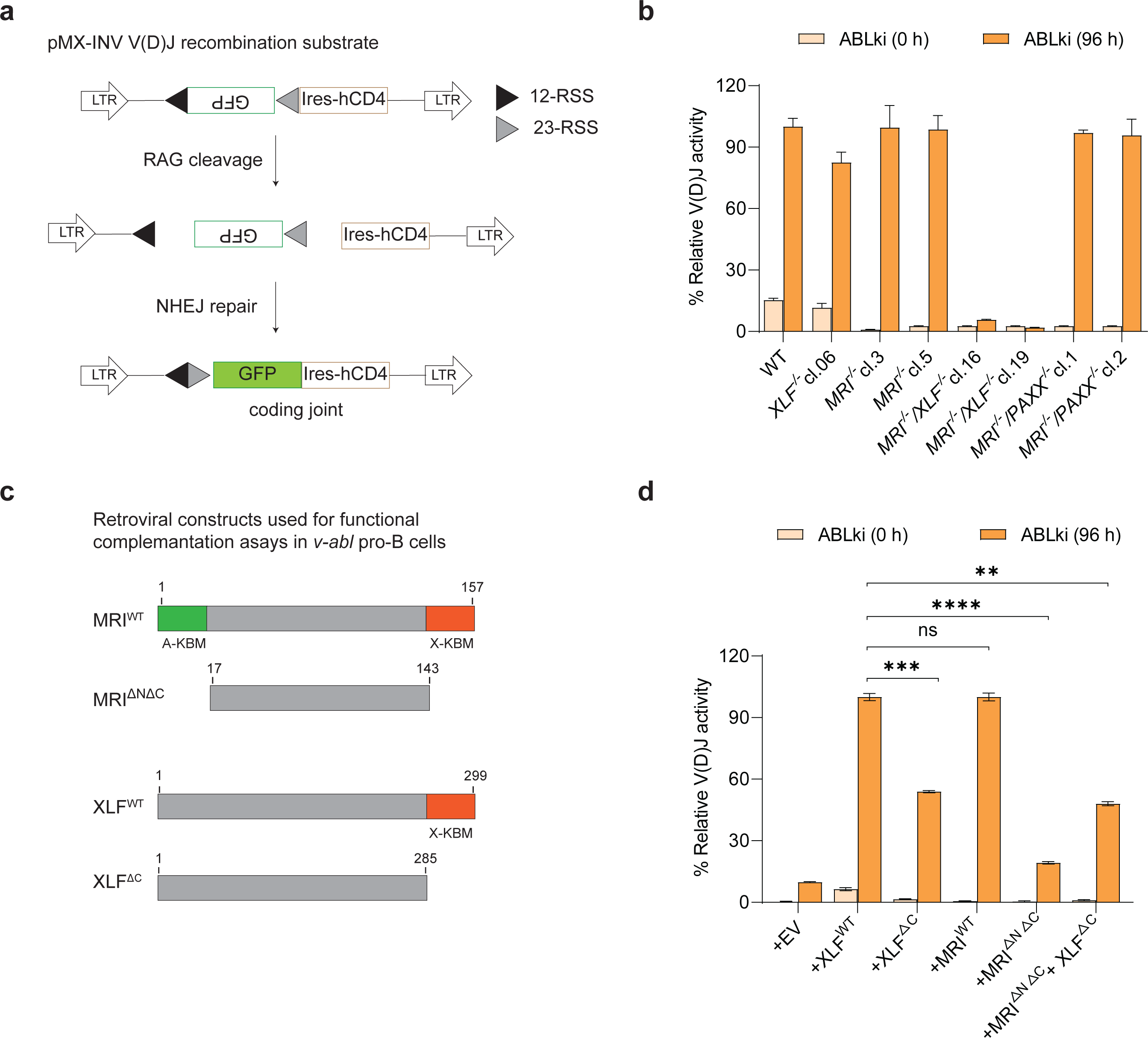
Combined deletion of the KBMs from MRI and XLF blocks NHEJ-mediated repair of RAG-DSBs. **a** Schematic representation of pMX-INV recombination substrate, following RAG cleavage, and the resulting coding joint (CJ) product by NHEJ. The long-terminal repeats (LTR), GFP cDNA, recombination signals (RSS, represented by 12-RSS; black triangle, 23-RSS; grey triangle), IRES-human CD4 cDNA (IRES-hCD4) are shown. **b** Percentage of relative V(D)J activity in WT, *XLF^-/-^* and 2 independent clones each from *MRI^-/-^*, *MRI^-/-^/XLF^-/-^*, and *MRI^-/-^/PAXX^-/-^ v-abl* pro-B cell clones treated for 0, and 96 h with ABLki were assayed for pMX-INV rearrangement by flow cytometry. **c** Scheme shows MRI^WT^, MRI^ΔNΔC^, XLF^WT^ and XLF^ΔC^ variant retroviral constructs used in the functional complementation assays. The A-KBM (green box) and X-KBM (red box) are indicated. **d** Flow cytometric analyses relative V(D)J activity in *MRI^-/-^/XLF^-/-^ v-abl* pro-B cells retrovirally transduced with empty vector (EV), XLF^WT^, XLF^ΔC^, MRI^WT^, MRI^ΔNΔC^, and combination of MRI^ΔNΔC^+ XLF^ΔC^ treated with ABLki for 0 and 96 h. Histograms represent means (± SD) of at least three independent experiments of transduced cells in MRI*^-/-^/XLF^-/-^ v-abl* pro-B cells.

### Combined deletion of KBMs in PAXX and XLF failed to restore V(D)J activity

We and others previously established the functional redundancy between XLF and PAXX, resulting in synthetic lethality in double deficient mice and impaired V(D)J recombination both *in vivo* and *in vitro* ^27, 28, 30^. This prompted us to further investigate the role of XLF and PAXX C-terminal KBMs in CRISPR/Cas9 generated *PAXX^-/-^* and *PAXX^-/-^/XLF^-/-^ v-abl* pro-B cells (Fig. S2a, S2b). ABLki treatment of *PAXX^-/-^ v-abl* pro-B cells containing pMX-INV resulted in V(D)J recombination at similar levels as WT *v-abl* pro-B cells, in accordance with the absence of major immune phenotype in *PAXX* KO mice (Fig. 2a). In contrast and as expected, V(D)J recombination was severely impaired in *PAXX^-/-^/XLF^-/-^ v-abl* pro-B cells (Fig. 2a). To substantiate the need for KBMs in PAXX and XLF during RAG DSB repair, V(D)J recombination was assayed in *PAXX^-/-^/XLF^-/-^* cells by complementing with XLF^WT^, PAXX^WT^ or with XLF^ΔC^ and PAXX^ΔC^ variants, lacking their known respective KBMs (panel below Fig. 2b). Introduction of XLF^WT^ or PAXX^WT^ restored V(D)J recombination of MX-INV recombination substrate in *PAXX^-/-^/XLF^-/-^ v-abl* pro-B cells, as indicated by GFP expression after ABLki treatment (Fig. 2b). This functional complementation was dependent upon the presence of the respective KBMs as shown by the decreased activity when using XLF^ΔC^ or PAXX^ΔC^. Moreover, the co-expression of the two factors lacking their respective KBMs (PAXX^ΔC^+ XLF^ΔC^) failed to fully complement the V(D)J activity, thus further strengthening our hypothesis that the redundancy between PAXX and XLF, similarly to MRI and XLF, is likely controlled by their interaction with Ku for the repair of RAG induced DSBs during V(D)J recombination (Fig. 2b).

**Fig. 2.**
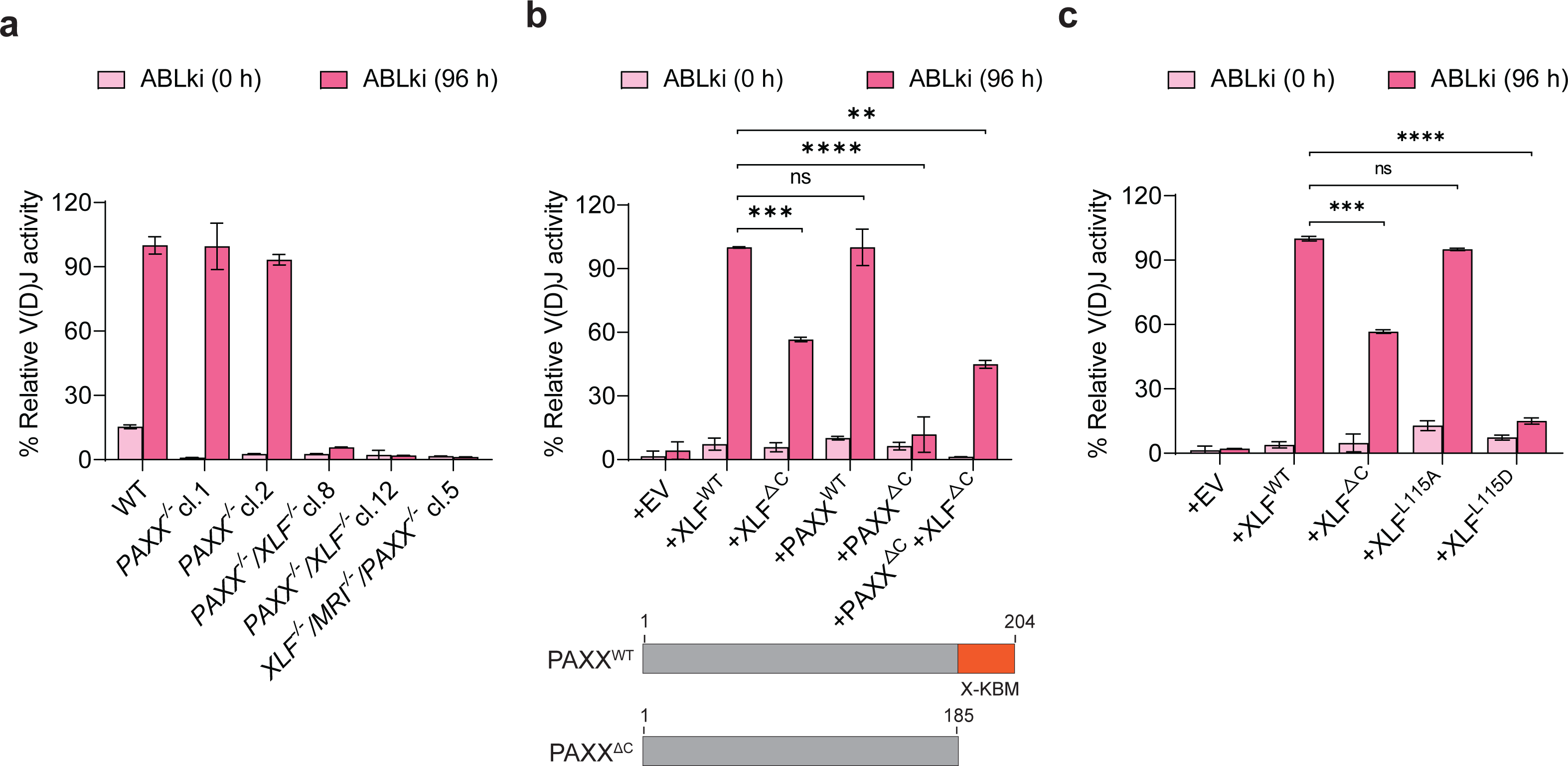
Combined deletion of the KBMs from PAXX and XLF weakens NHEJ-mediated repair of RAG-DSBs. **a** Relative V(D)J activity in WT, 2 independent clones each from *PAXX^-/-^, PAXX^-/-^/XLF^-/-^,* and MRI*^-/-^/PAXX^-/-^/XLF^-/-^ v-abl* pro-B cell clones treated for 0, and 96 h with ABLki were assayed for pMX-INV rearrangement by flow cytometry. **b** Flow cytometric analyses of relative V(D)J activity in *PAXX^-/-^/XLF^-/-^* abl pro-B cells retrovirally transduced with EV, XLF^WT^, XLF^ΔC^, PAXX^WT^, PAXX^ΔC^, and combination of PAXX^ΔC^+XLF^ΔC^ treated with ABLki for 0 and 96 h. Panel below shows schematic representation of retroviral constructs used for functional complementation assays. The X-KBM (red box) is indicated. **c** Flow cytometric analyses of relative V(D)J activity in *XLF^-^*^/-^*/MRI^-/-^/PAXX^-/-^ v-abl* pro-B cells retrovirally transduced with EV, XLF^WT^, XLF^ΔC^, XLF^L115A^ and XLF^L115D^ treated with ABLki for 0 and 96 h. Histograms represent means (± SD) of at least three independent experiments.

Overall, expression of XLF^ΔC^ in XLF deficiency backgrounds, *MRI^-/-^/XLF^-/-^* (Fig. 1d) and *PAXX^-/-/^XLF^-/-^* (Fig. 2b), resulted in V(D)J activity greater than that provided by KBMs-deficient MRI or PAXX. We hypothesized that this residual V(D)J activity could be due to the presence of PAXX in *MRI^-/-^/XLF^-/-^*or MRI in *PAXX^-/-^/XLF^-/-^* respectively, suggesting some combinatorial redundancy among these various KBM-containing factors. To further investigate this idea, an *XLF^-/-^/MRI^-/-^/PAXX^-/-^*triple knockout *v-abl* pro-B clone was generated using CRISPR/Cas9 (Fig. S2b). V(D)J recombination was assayed in this clone upon transduction of retroviral constructs expressing EV (empty vector), XLF^WT^, and XLF^ΔC^, variants. As expected, complementation with XLF^WT^, restored V(D)J recombination, as indicated by GFP expression after ABLki treatment (Fig. 2c), while complementation with XLF^ΔC^ again restored V(D)J activity only to some extent.

Although XLF^ΔC^ lacks its interaction with Ku, it might still interact with XRCC4 (X4) or bind DNA on its own to partly stabilize the RAG induced DSBs. To abolish the interaction between XLF and X4, XLF^L115A^ and XLF^L115D^ mutants were generated. L115 residue of XLF is critical for interaction with X4 ^42, 43^. *In vitro* biochemical studies showed that XLF^L115A^ substitution does not interact with X4 ^42, 43^ but is still able to stimulate DNA ligation during NHEJ *in cellulo* ^44^ while XLF^L115D^ substitution, which also alters interaction with X4, failed to reverse the NHEJ deficits or radiosensitivity associated with XLF deficiency in fibroblasts ^43^. In agreement, complementation with XLF^L115A^ restored V(D)J activity in *XLF^-/-^/MRI^-/-^/PAXX^-/-^*cells similar to WT cells while cells complemented with XLF^L115D^ remained V(D)J recombination deficient (Fig. 2c), suggesting that in addition to *bona fide* XLF/Ku interaction, the capacity of XLF to stimulate the X4-LIG4 complex also contributes to DSBs repair during VDJ recombination.

### X-KBM of XLF is required for efficient RAG DSB repair

Then, to analyze the role of KBMs present in NHEJ factors for functional complementation with RAG2, a series of *v-abl* cells with disrupted *XLF*, *MRI* and *PAXX* genes in *RAG2^c/c^*background were generated using CRISPR-Cas9, resulting in *RAG2^c/c^/XLF^-/-^, RAG2^c/c^/MRI^-/-^, RAG2^c/c^/PAXX^-/-^* genotypes (Fig. S3a-S3c). *RAG2^c/c^/XLF^-/-^* cells exhibited complete block in V(D)J recombination (Fig. 3a) owing to the abolition of the functional redundancy between R2CT and XLF for the synapsis of RAG induced DSBs ^23^. In contrast, RAG*2^c/c^/MRI^-/-^*and *RAG2^c/c^/PAXX^-/-^* cells showed no detectable V(D)J recombination defect, arguing that R2CT is epistatic with MRI and PAXX, as previously reported for PAXX ^26, 27^. To further investigate the function of XLF KBM as part of the functional redundancy with R2CT during V(D)J recombination, functional complementation of *RAG2^c/c^/XLF^-/-^* cells with retroviral constructs expressing EV, XLF^WT^, and XLF^Δ^ were performed. The V(D)J activity of *RAG2^c/c^/XLF^-/-^* cells complemented with XLF^ΔC^, lacking the X-KBM motif, was only half of that observed with full length XLF^WT^ (Fig. 3b, 3c), in agreement with the lack of complementation by XLF^ΔC^ in the XLF deficiency backgrounds *MRI^-/-^/XLF^-/-^* (Fig. 1d) and *PAXX^-/-^/XLF^-/-^* (Fig. 2b). Altogether, these data support that the X-KBM of XLF and c-terminus of RAG2 participate in the functional redundancy of the two factors during the NHEJ-mediated repair of RAG-DSBs to maintain DNA end synapsis, ensuring efficient DNA repair in G1 phase lymphocytes.

**Fig. 3.**
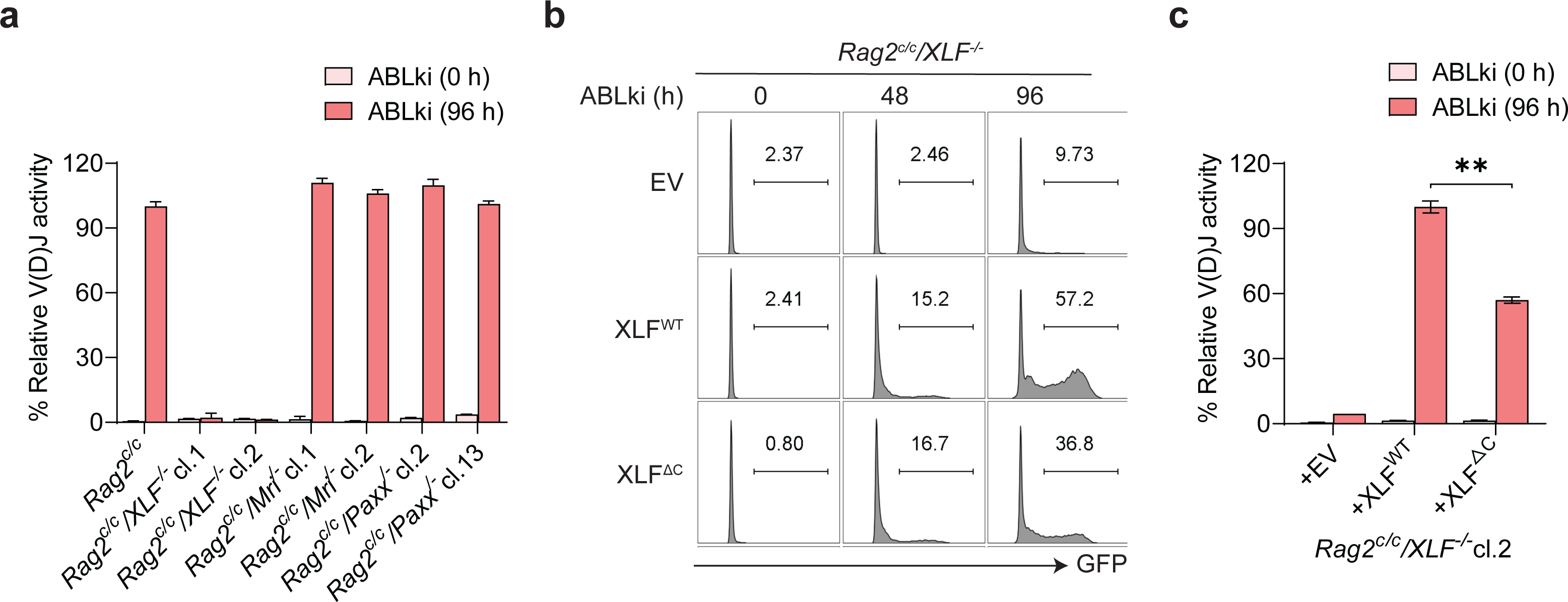
X-KBM of XLF is required for its function during V(D)J recombination. **a** *RAG2^c/c^*, *RAG2^c/c^/XLF^-/-^*, *RAG2^c/c^/MRI^-/-^*, *RAG2^c/c^/PAXX^-/-^ v-abl* pro-B cell clones (2 independent clones each) treated for 0, and 96 h with ABLki were assayed for pMX-INV rearrangement by flow cytometry, with the percentage of relative V(D)J activity indicated. **b, c** Flow cytometric analyses of GFP expression and the percentage of relative V(D)J activity respectively in *RAG2^c/c^/XLF^-/-^ v-abl* pro-B cells retrovirally transduced with EV, XLF^WT^, and XLF^ΔC^ and treated with ABLki for the indicated lengths of time. Histograms represent means (± SD) of at least three independent experiments of transduced cells in *RAG2^-/-^/XLF^-/-^ v-abl* pro-B cell.

### Identification of a KBM in the RAG2 C-terminal Region

Given the critical role of the KBM from XLF for RAG2/XLF functional redundancy, as for MRI and PAXX, we asked whether this could also be true for RAG2. Multiple sequence alignment of metazoans RAG2 sequences allowed to identify a conserved region in the last 39 amino acids of R2CT (Fig. 4a). This region can be divided in two main motifs rich in basic residues, which interestingly resemble KBM motifs of several DNA repair factors (Fig. 4b). An extreme c-terminal Motif 2 aligns well with X-KBM and P-KBM present in XLF and PAXX, respectively. The aspartic acid located at the end of the motif would be more in favor of a P-KBM, while X-KBM harbors a serine at this position. An upstream Motif 1, although rich in basic residues, only loosely aligns with known KBMs but shows striking conservation with a short peptide in PAXX also present just upstream of the *bona fide* P-KBM. To further document the reality of a KBM in the R2CT, we evaluated the interaction between Ku and peptides derived from this region through Isothermal Titration Calorimetry (ITC) (Fig. 4c-e, S4), the results of which are summarized in (Table 1). Ku70-Ku80 deleted from its flexible C terminal regions (called Ku^ΔC^) were purified similar to our previous studies with APLF, XLF and PAXX-KBM (Fig. S4a) ^32, 37, 45^. The interaction of the RAG2 Motifs 1 and 2 with Ku^ΔC^ bound to a 42 bp DNA was measured as previously done for PAXX-KBM ^45^ and no direct interaction was observed with either motifs alone motif 1 (RAG2a) and motif 2 (RAG2b), (Table 1, lines 1, 2, Fig. S4b, S4c). In contrast, a micromolar interaction appeared with the longer peptide containing both motifs (K_D_ of 0,84 +/- 0,05 µM) (Table 1, line 3, Fig. 4d). This indicates that both motifs 1 & 2 are necessary for the interaction and that the resulting long c-terminal sequence indeed constitutes a new KBM named RAG2-KBM or R-KBM. In parallel, the interaction of this newly defined R-KBM with purified full-length Ku70/Ku80 heterodimer (Fig. S4a, Table 1, line 4, S4d) was further characterized. The full-length heterodimer revealed a similar affinity to Ku^ΔC^ suggesting the c-terminal regions of Ku are not involved in the interaction with R-KBM (K_D_ of 1,18 +/- 0,19 µM). Next, to evaluate if the interaction of R-KBM with Ku^ΔC^ is DNA dependent, the interaction of R-KBM with Ku^ΔC^ was measured in the absence of DNA and no interaction was observed (Table 1, line 5, Fig. S4e). These observations are similar to PAXX-KBM interactions from our previous report ^45^ but different with APLF- and XLF-KBM that interact similarly with or without DNA ^37^. We then tested the interaction with Ku^ΔC^ bound to a DNA containing a 15 bp duplex part and a 15 nt 5’ -overhang as done previously with the PAXX-KBM. The affinity is ∼7 fold weaker than with the 42 bp DNA (Table 1, line 6, compare Fig. S4f, 4f). This differs from PAXX-KBM that presented similar affinity with the two different DNA ^45^. The sequence alignment of R-KBM with the other known KBMs shows some sequence similarity with the PAXX-KBM (Fig. 4b). To further assessed possibility of a competition between the PAXX-KBM and the R-KBM, we titrated with the R-KBM a Ku^ΔC^-DNA (42 bp) complex incubated with an excess of PAXX-KBM. We observed an interaction with a marginally weaker affinity suggesting an absence of competition under ITC conditions (Table 1, line 7, Fig. S4g). Lastly, the interactions with Ku of three mutants of the R-KBM, either in motif 1 (conserved K503 to E) or motif 2 (conserved F526 to E or double mutant of conserved L525-F526 to E) were measured (Table 1, lines 8-10, Fig. 4e, S4h, S4i). We observed no interaction with the RAG2-KBM K503E and a ∼1.5-2-fold reduction of affinity with the F526E and L525E-F526E mutants. As control, the heat exchange was measured following RAG2c injection into a micro-calorimeter cell containing the buffer, and we observed no heat exchange during these experiments (Fig. S4j). Altogether, these results confirm that the last 39 residues of RAG2 define a new KBM that binds Ku *in vitro* in a DNA-dependent manner.

**Fig. 4.**
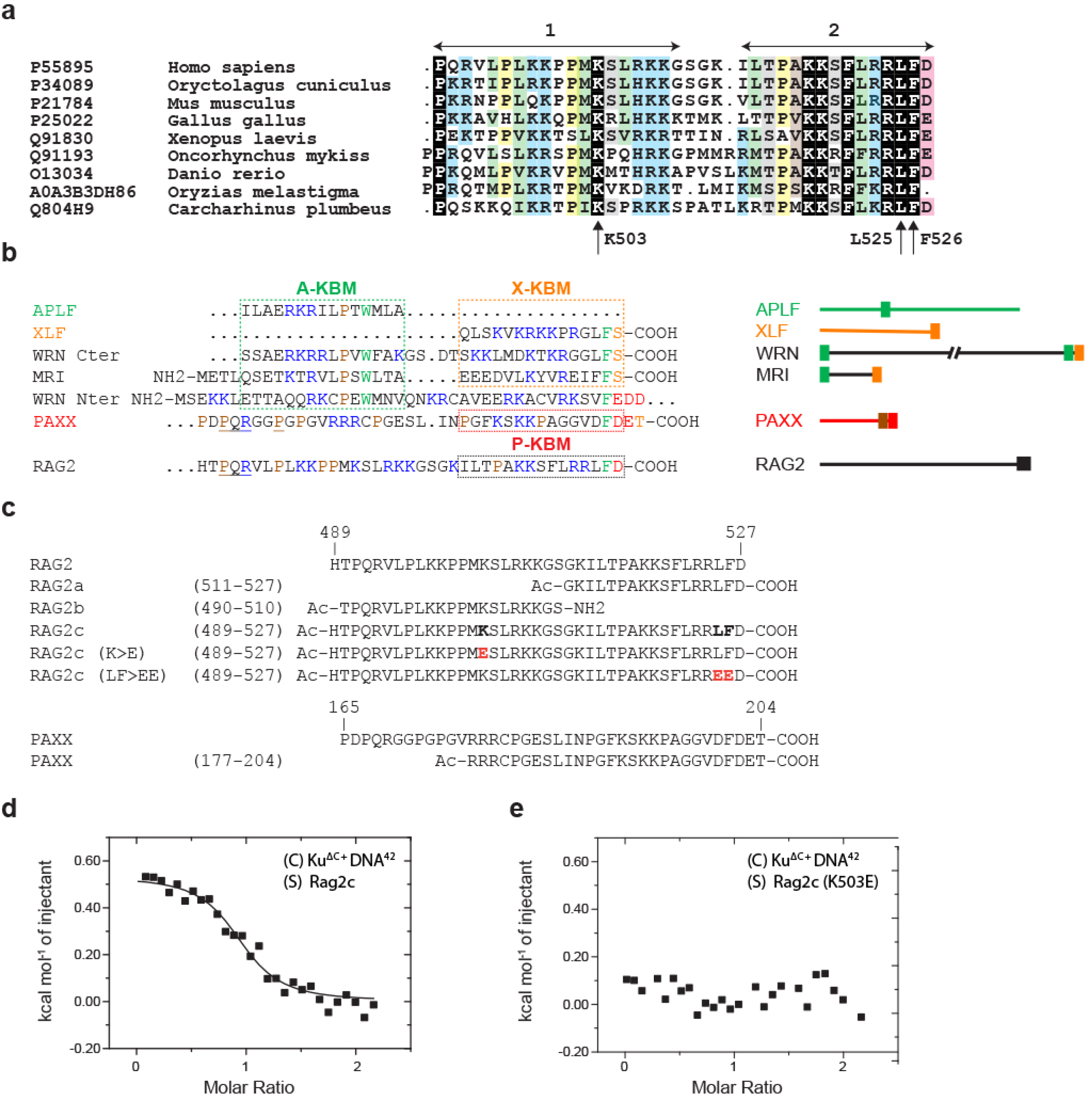
Identification of a KBM in the RAG2 C-terminal region. **a** Alignment of the amino acids sequences of the RAG2 C-terminal in various species, from human to sandbar shark (UniProt identifiers are given in front of the species names). **b** Alignment of known KBM amino acid sequences of APLF, XLF, MRI, WRN, PAXX and RAG2. Ku binding motifs (KBM) are classified according to their representative leader (A- for APLF, X- for XLF and P- for PAXX). Basic (blue), acidic (red), aromatic (green) and polar (orange) residues are highlighted. KBMs are positioned on the corresponding protein primary sequences (right). **c** Sequences of the peptides used for ITC measurements. Rag2a and Rag2b are respectively motifs 1 and 2, Rag2c contains both motifs and constitutes the R-KBM. Rag2c (K>E) and Rag2c(LF>EE) are two mutants respectively in motifs 1 and 2. PAXX(177-204) is the peptide containing the P-KBM. The sequence above these peptides are the sequence of the proteins. The N-ter of the peptides are acetylated to mimic the peptide bond. The c-terminus of the peptide are carboxylate when it is the last residue of the protein or it is amidated if it is an internal position to mimic the peptide bond. (d) Isotherm of titration of Ku^ΔC^-DNA^42^ complex titrated by peptide Rag2c obtained by ITC. The association constant Ka and the thermodynamic parameters measured are reported in Table1 (lane 3). Ku^ΔC^ is a Ku construct deleted of its c-terminal region and DNA^42^ is a 42bp DNA duplex (e) Isotherm of titration of Ku^ΔC^-DNA^42^ complex titrated by peptide Rag2c (K>E), mutant K503E in motif 1 (see Table 1, lane 10).

**Table 1.** Characterization of RAG2 c-terminal region interactions with Ku-DNA complexes by Micro-calorimetry.

| N° | ITC Cell | ITC Syringe | $K_a$ ($10^5$ M) | $K_d$ (μM) | $\Delta H^\circ$ (kcal/mol) | $-T\Delta S^\circ$ (kcal/mol) | $\Delta G^\circ$ (kcal/mol) | Observation | Thermogram |
| --- | --- | --- | --- | --- | --- | --- | --- | --- | --- |
| 1 | Ku<sup>ΔC</sup>-DNA<sub>42</sub> 1:1.1 | RAG2a | NI | NI | NI | NI | NI | Peptide residues 511-527, GKILTPAKKSFLRRLFD | Fig. S4b |
| 2 | Ku<sup>ΔC</sup>-DNA<sub>42</sub> 1:1.1 | RAG2b | NI | NI | NI | NI | NI | Peptide residues 490-510, TPQRVLPLKKPPMKSLRKKGS-NH2 | Fig. S4c |
| 3 | Ku<sup>ΔC</sup>-DNA<sub>42</sub> 1:1.1 | RAG2c | 12.0 ± 0.6 | 0.84 ± 0.05 | 0.525 ± 0.009 | -8.81 ± 0.04 | -8.29 ± 0.03 | Peptide residues 489-527, HTPQRVLPLKKPPMKSLRKKGSGKILTPAKKSFLRRLFD | Figure 4f |
| 4 | Ku<sup>FL</sup>-DNA<sub>42</sub> 1:1.1 | RAG2c | 8.7 ± 1.4 | 1.2 ± 0.2 | 0.5 ± 0.1 | -8.55 ± 0.03 | -8.1 ± 0.1 | No effect of the Cter K70 and Cter Ku80 on the affinity | Fig. S4d |
| 5 | Ku<sup>ΔC</sup> | RAG2c | NI | NI | NI | NI | NI | RAG2c versus Ku heterodimer without DNA | Fig. S4e |
| 6* | Ku<sup>ΔC</sup> – DNA<sub>ov</sub><br>(15 bp–15 ss) 1:1.1 | RAG2c | 1.7 ± 0.4 | 5.8 ± 1.3 | 1.5 ± 0.1 | -8.7 ± 0.3 | -7.2. ± 0.1 | 5' ssDNA extension | Fig. S4f |
| 7 | Ku<sup>ΔC</sup> - DNA<sub>42</sub>-PAXX<br>1:1.1:10 | RAG2c | 5.1 ± 1.7 | 2.2 ± 0.7 | 0.8 ± 0.3 | -8.6 ± 0.1 | -7.8 ± 0.2 | No competition with PAXX peptide residues 178-204, RRRCPGESLINPGFKSKKPAGGVDFDET | Fig. S4g |
| 8 | Ku<sup>ΔC</sup>-DNA<sub>42</sub> 1:1.1 | RAG2c (F>E) | 8.1 ± 4.4 | 1.7 ± 0.9 | 0.9 ± 0.3 | -8.9 ± 0.1 | -8.0 ± 0.4 | No loss of interaction between Ku heterodimer and RAG2c mutant | Fig. S4h |
| 9* | Ku<sup>ΔC</sup>-DNA<sub>42</sub> 1:1.1 | RAG2c (LF>EE) | 6.7 ± 1.1 | 1.5 ± 0.3 | 0.92 ± 0.03 | -8.9 ± 0.1 | -8.0 ± 0.1 | No loss of interaction between Ku heterodimer and RAG2c mutant | Fig. S4i |
| 10 | Ku<sup>ΔC</sup>-DNA<sub>42</sub> 1:1.1 | RAG2c (K>E) | NI | NI | NI | NI | NI | Loss of interaction between Ku heterodimer and RAG2c mutant | Fig. 4g |
| 11 | RAG2c | Buffer | No signal |  |  |  |  |  | Fig. S4j |

Representative thermograms are reported in Figure XXXX. All ITC experiments in this study were realized at 25C. NI means no interaction, small and constant signal due to the dilution of the molecules injected in the cell. The error is the deviation to the mean value of at least two measurements.

### The RAG2 C-terminal region forms a stable complex with Ku

To further validate the identification of R-KBM in R2CT, we interrogated unbiased R2CT-interactome in *v-abl* pro-B cells. *v-abl* pro-B cells were transduced with MSCV-GFP-R2CT or MSCV-GFP retroviral supernatants to stably express either a N-terminal GFP-tagged R2CT (aa351-527) fusion protein or GFP alone as a control, respectively (Fig. S5a). The R2CT contains a cell cycle regulated protein degradation signal in which the phosphorylation of an essential threonine (T490 in full-length RAG2) signals the periodic destruction of RAG2 at the G1-to-S transition and therefore ensures proper coupling of V(D)J recombination to the cell cycle ^46, 47^. Consistent with these findings, flow cytometry analysis showed that, in contrast to *v-abl* pro-B cells transduced with MSCV-GFP, cycling *v-abl* pro-B cells transduced with MSCV-GFP-R2CT expressed poorly the GFP-R2CT protein in the absence of ABLki (0 h) (∼33% GFP positive cells vs 93.2% in MSCV-GFP transduced cells), but reached up to 98% GFP positive cells upon treating the cells for 24, 48 and 72 h with ABLki, attesting for the time-dependent stabilization of GFP-R2CT in G0/G1 arrested cells (Fig. S5b). We interrogated the R2CT interactome after treating the *v-abl* pro-B cells for 72 h with ABLki, which represents the ideal physiological condition to identify R2CT interacting partners during the V(D)J recombination process. Protein complexes were recovered using nanobodies directed against the GFP moiety covalently linked to magnetic beads ^45, 48^ and analyzed by SDS-PAGE, which revealed the presence of several specific bands that were absent from the GFP alone pulldown (Fig. 5a, lanes 1, 2). Mass-spectrometry analyses of these complexes identified XRCC6 (Ku70) and XRCC5 (Ku80) among the top 30 hits (ProteomeXchange: PXD045640). Accordingly, endogenous Ku70 and Ku80 were efficiently and specifically recovered with GFP-R2CT (Fig. 5b). In order to assess the effect that contaminating DNA could have on GFP-R2CT complex recovery, samples were also treated with ethidium bromide (EtBr) or Benzonase before elution. EtBr led to the disappearance of several prominent protein bands (Fig. 5a, compare lane 2, 4). Mass spectrometry analyses demonstrated that GFP-R2CT-Ku interaction was resistant to both EtBr and Benzonase treatments (Fig. 5c-5e). Thus, under the overexpression conditions in cells used here, GFP-R2CT associates with Ku and the interaction is largely DNA independent, although DNA was required to obtain measurable interaction *in vitro* between only RAG2 c-terminal peptide and purified Ku (see above).

**Fig. 5.**
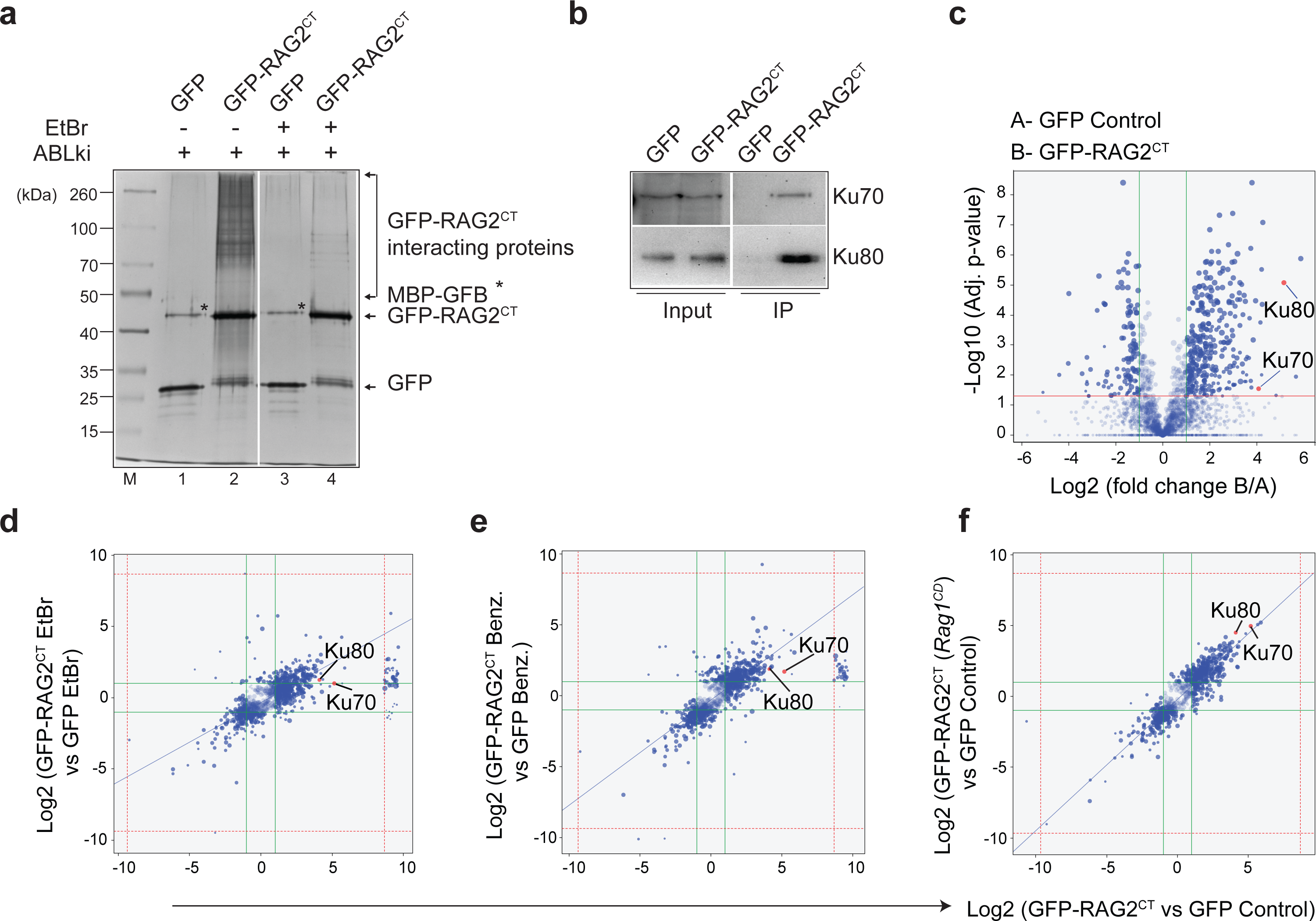
RAG2 C-terminal region interacts with Ku. **a** Coomassie Blue-stained reducing SDS-PAGE showing the polypeptides that were recovered in the presence or absence of Ethidium Bromide (EtBr) from extracts of *v-abl* pro-B cells expressing GFP-RAG2^CT^ or GFP as control. Cells were treated with 0.5 µM Abelson kinase inhibitor (ABLki) for 72 hours before harvesting. **b** Western blotting confirming specific recovery of Ku70 and Ku80 in the GFP-RAG2^CT^ pull-down samples. M = size markers. **c** Volcano plots comparing the total numbers of GFP-RAG2^CT^-associated peptides in WT *v-abl* pre-B cells. **d-f** Correlation plots comparing the total numbers of GFP-RAG2^CT^-associated peptides in the presence of Ethidium Bromide (EtBr), Benzonase (Benz), and from *RAG1* catalytic dead (*RAG1^CD^*) *v-abl* pro-B cells as determined by mass spectrometry.

### Interaction of RAG2 C-terminal region with Ku does not depend on RAG1/2-induced DSBs

To decipher whether GFP-R2CT-Ku interaction depends on the induction of RAG-DSBs, we utilized CRSPR-Cas9 generated *RAG1* catalytic dead (*RAG1^CD^*) *v-abl* pro-B cells that express GFP-R2CT (Fig. S6a). RAG1 contains RSS-binding domains, a region that interacts with RAG2, and an active site for DNA cleavage, which includes three essential acidic amino acids called DDE motif (D600, D708 and E962) that coordinate divalent metal ions and are essential for catalysis ^18, 49, 50^. CRISPR gRNAs were designed targeting the DDE regions and allowed to obtain *RAG1^CD^ v-abl* cells harboring an in-frame deletion of the highly conserved C599 residue, adjacent to the D600 aspartate (Fig. S6a). Immunoblotting showed that *RAG1^CD^* cells express RAG1 protein levels similar to the WT cells (Fig. S6b). Treatment of WT cells with ABLki triggered *Vk*-to-*Jk* rearrangement at the endogenous *Igk* locus, as evidence by PCR analysis of inversional *Igk* V_6–23_-*J1* revealing CJ formation (Fig. S6c) while no such CJ could be PCR amplified in *RAG1^CD^* cells under similar conditions of RAG induction. These results attest for the RAG1 loss of function in *RAG1^CD^* cells. In order to check whether the R2CT and Ku interaction depends on RAG1 activity, R2CT interactome was performed using *RAG1^CD^*cells. Mass-spectrometry analysis showed the presence of Ku70/80 in GFP-R2CT precipitates to the same extent as in WT cells suggesting the interaction between Ku and R2CT is independent of RAG1 activity (Fig. 5f). To further strengthen these observations, we performed Proximity Ligation Assay (PLA), which enables visualization and quantification of protein interactions *in cellulo*. *v-abl* pro-B cells expressing GFP-R2CT or GFP alone were stained with antibodies (Abs) against different combinations of GFP and Ku and subsequently with secondary Abs conjugated with oligonucleotides (PLA probes) to generate a strong red fluorescence dot when the two PLA probe–marked proteins are in close proximity (≤40 nm) ^51, 52^. As shown in (Fig. 6a, b), similar PLA-signals were obtained for GFP and Ku80 in the WT and *RAG1^CD^ v-abl* pro-B cells expressing GFP-R2CT, comparable to the positive control for Ku70/Ku80 PLA. In contrast, the PLA-signal was minimal in the control cells expressing GFP alone, as well as when one or both of the primary Abs were omitted. Taken together, these data establish the existence of GFP-R2CT-Ku complex formation *in cellulo*, an interaction that is largely independent of RAG1 activity.

**Fig. 6.**
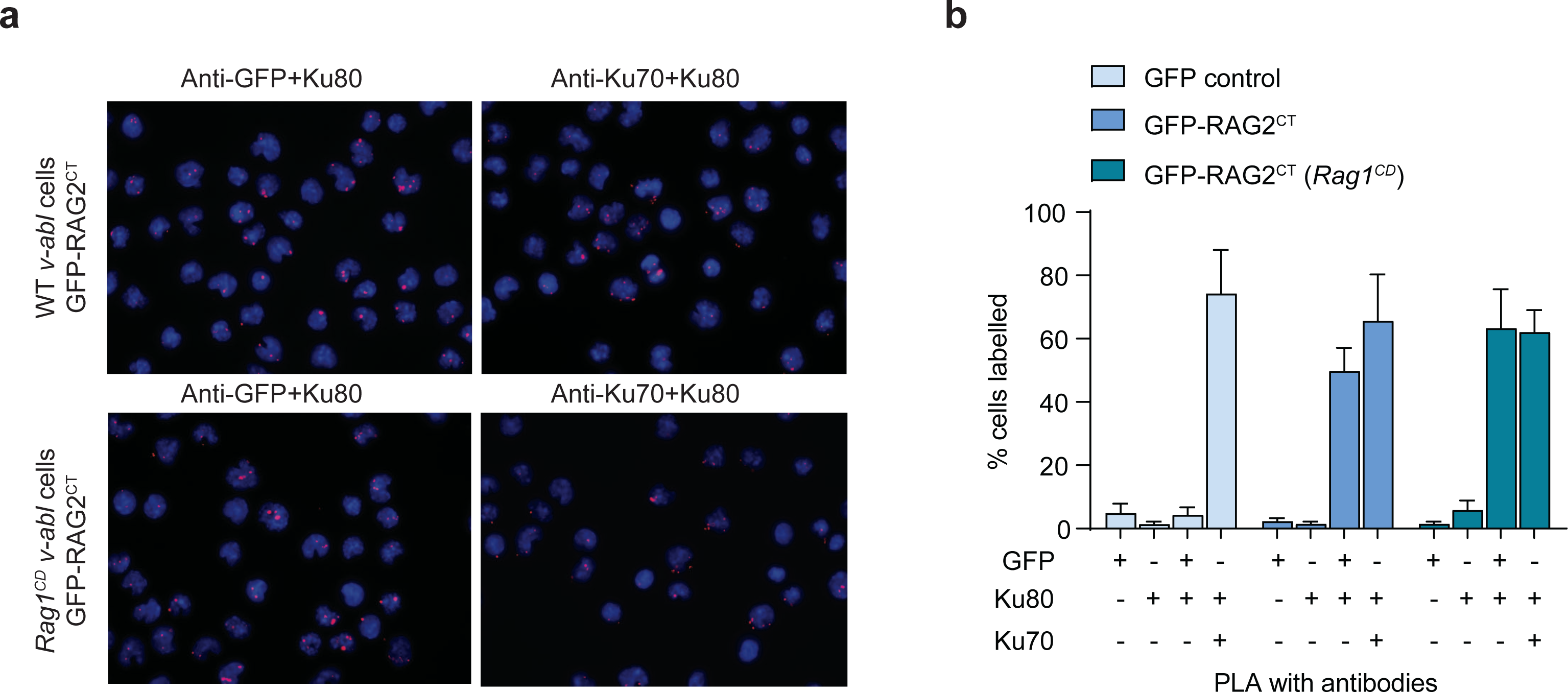
*In situ* PLA analysis to visualize the interaction between RAG2 c-terminal region and Ku complex. **a** Detection of the proximal location of each protein pair analyzed (visualized as red dots) by in situ PLA conducted on ABLki-treated WT and *RAG1^CD^ v-abl* pro-B cells (original magnification 63X). **b** Summary of the PLA results showing the percentage of cells labelled for the indicated protein pairs. Number of labelled cells were calculated relative to those of cells controls (cells expressing GFP alone, as well as when one of the primary Abs were omitted. Data were quantified from three independent experiments where error bars are indicated (± SD).

### Modelling RAG2 C-terminal region and Ku interaction through AlphaFold2

We used AlphaFold2-multimer, run on ColabFold v1.5.2 ^53^ to model the interaction of the R-KBM (aa 481-527) with Ku. Using first full Ku70/80 heterodimer, this led to identify Ku80 as the target of RAG2, with a common anchor point observed for the 5 proposed models at the position of the A-KBM binding site on the vWA domain. Then we used the same RAG2 peptide and only the vWA domain of Ku80 (aa 1-245), in order to refine the prediction (Fig. S7 a-c). Two anchor points were consistently observed for the five proposed models, at the positions of the A-KBM binding site for RAG2 motif 1 and of the X-KBM binding site for RAG2 motif 2. These anchor points are separated by a large, disordered linker (Fig. 7a). The RAG2 C-terminal end (motif 2) fits well into the X-KBM binding site, superimposing on the XLF peptide, as observed in the experimental 3D structure of the complex (pdb 6ERG) (Fig. 7b, c) and consistent with the sequence alignment between RAG2 and XLF (Fig. 4b). Despite very low pLDDT, the predicted alpha helical conformation is supported by the hydrophobic amino acid periodicity, and the conformations observed for other X-KBM sequences. The RAG2 N-terminal end (motif 1) fits into the A-KBM binding site. Here, even though all five models propose binding in this site, there is a much larger variety of conformations, again with very low pLDDT, no clear superimposition with the APLF peptide, as observed in the experimental 3D structure of the complex (pdb 6ERF) and no obvious correspondence with the proposed alignment of the two sequences (Fig. 4b). Nevertheless, further investigations at the experimental level are now needed to clarify the specific details of the RAG2-Ku interaction at the atomic level.

**Fig. 7.**
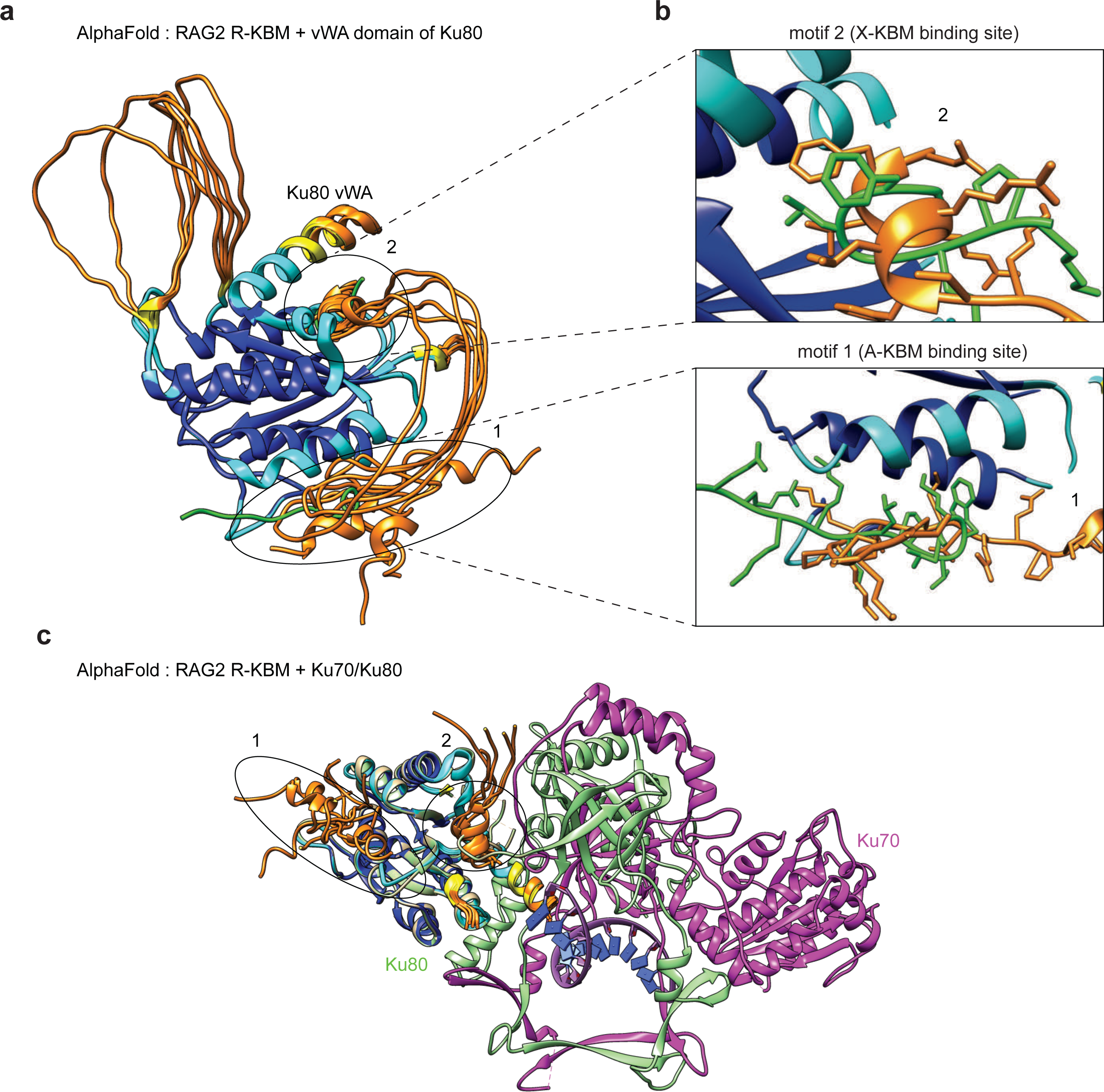
AlphaFold2 (AF2) model of R2CT/Ku interaction. **a** Superimposition of the five AF2 3D structure models of the RAG2 extended KBM (aa 481-527) in complex the vWA domain of human Ku80 (aa 1-245). These were superimposed with the experimental 3D structures of Ku80 in complex with XLF (pdb 6ERG) and Ku80 in complex with APLF (pdb 6ERF) (only the XLF and APLF peptides are shown in green, at the level of the N-terminal (motif 1) and C-terminal (motif 2) regions of the RAG2 peptides). The AF2 models are colored according to the pLDDT values (orange: very low (<50), yellow: low (between 50 and 70), high (between 70 and 90), very high (>90). **b** Focus on motifs 1 (A-KBM binding site) and 2 (X-KBM binding site). **c** Superimposition of the 3D structure models of the RAG2 extended KBM in complex the vWA domain of human Ku80 onto the Ku70/Ku80 complex (pdb 6ERF). For clarity, the RAG2 central disordered linker and the Ku80 disordered loop (aa 170-195) are not shown.

### NHEJ function of RAG2 C-terminal region depends on the R-KBM

Lastly, to determine whether R-KBM functions in the repair of RAG-mediated DSBs, *RAG2^-/-^* and *RAG2^-/-^/XLF^-/-^ v-abl* pro-B cell clones were generated through CRISPR-Cas9 for complementation studies (Fig. S8a). Flow cytometry analyses in *v-abl* pro-B cells with WT, *XLF^-/-^*, *RAG2^-/-^*, *RAG2^-/-^/XLF^-/-^* genotypes showed robust levels of pMX-INV rearrangement in 96 h ABLki-treated WT and *XLF*^-/-^ cells as expected, with 64% and 55% GFP positive cells respectively (Fig. 8a). In contrast, no GFP positive cells were recovered from ABLki treated *RAG2^-/-^* and *RAG2^-/-^/XLF^-/-^*cells, confirming the expected lack of V(D)J recombination in the context of a *RAG2* loss of function (Fig. 8a and Fig. S8b). To test explicitly the involvement of RAG2 KBMs during V(D)J recombination, functional complementation studies were performed in *RAG2^-/-^* and *RAG2^-/-^*/*XLF^-/-^*cells with retroviral constructs expressing EV, RAG2^WT^, RAG2^Δ1^ and RAG2^Δ2^, lacking either motif 2 or both motifs 1 and 2 of the R-KBM, respectively (Fig. 8b). While expression of RAG2^WT^ and RAG2^Δ1^ led to full complementation of V(D)J activity in both *RAG2^-/-^* and *RAG2^-/-^*/*XLF^-/-^* cells (Fig. 8c, d), V(D)J complementation with RAG2^Δ2^ (lacking R-KBM) was impaired in these cells. These results suggest that, in the absence of XLF, the functional redundancy provided by the R2CT domain partly relies on the interaction with Ku via the R-KBM present in this region of RAG2. Nevertheless, the residual V(D)J activity provided by the RAG2^Δ2^ mutant in *RAG2^-/-^/XLF^-/-^* cells contrasts with the known impaired V(D)J activity in *RAG2^c/c^*/*XLF^-/-^*^23^ cells and suggests that other determinants present in R2CT outside R-KBM participate in the R2CT and XLF redundancy.

**Fig. 8.**
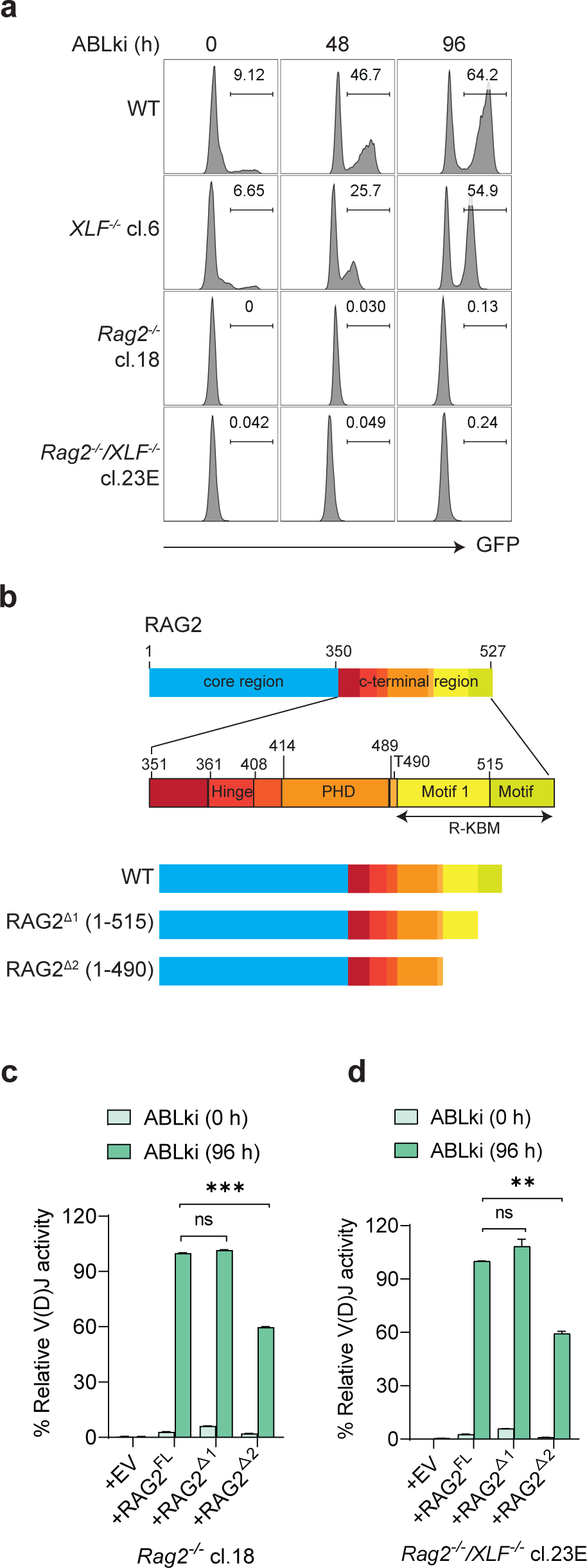
R-KBM of RAG2 is required during V(D)J recombination in *v-abl* pro-B cells. **a** Flow cytometric analysis of GFP expression in WT, *XLF^-/-^, RAG2^-/-^* and *RAG2^-/-^/XLF^-/-^ v-abl* pro-B cells treated with ABLki for the indicated lengths of time. **b** Scheme shows WT RAG2 full length protein highlighting the core region and the extreme c-terminal region with the identified motifs 1 and 2 and RAG2 variant constructs used in the study. **c, d** Flow cytometric analyses of percent relative V(D)J activity in *RAG2^-/-^* and *RAG2^-/-^ /XLF^-/-^* abl pro-B cells retrovirally transduced with empty vector EV, RAG2^FL^, RAG2^Δ1^ and RAG2^Δ2^ treated with ABLki for at 0 and 96 h. Histograms represent means (± SD) of at least three independent experiments of transduced cells in *RAG2^-/-^* and *RAG2^-/-^/XLF^-/-^ v-abl* pro-B cells.

## Discussion

DSBs are considered the most toxic DNA lesions. Nevertheless, programmed DSBs (prDSBs) are part of several physiological processes in the living kingdom. Perhaps the most spectacular instance of such prDSBs lie in the excision of thousands of Internal Eliminated Sequences (IES) from the genome of unicellular ciliates such as *Paramecium* during their life cycle through a precise “cut-and-close” mechanism by the domesticated transposase PiggyMAC ^54^. In higher eukaryotes, the diversification of the adaptive immune system in developing B and T lymphocytes is achieved, by the somatic DNA rearrangement of their Immunoglobulin (Ig) and T Cell receptor genes respectively through the mechanism of V(D)J recombination ^13–15^. This molecular process also proceeds through the introduction of prDSBs via the domesticated transposase RAG1/2. In these two examples, the introduction of prDSBs, although highly hazardous, appears mandatory for the viability of the *Paramecium* or the proper maturation of B and T lymphocytes. PiggyMAC deficient *Paramecium* are not viable ^55^ and RAG1/2 loss of function results in abortive lymphoid development, leading to severe combined immunodeficiency (SCID) in both humans and animal models ^56^. The NHEJ DNA repair pathway copes with the prDSBs efficiently in both instances. Nonetheless, one of the major threats posed by DSBs in general and prDSBs in particular is the possibility, in a context of DNA repair dysfunction, of having free DNA ends hanging around, a major subset for chromosomal translocation, rendering V(D)J recombination a potential oncogenic process. Accordingly, several engineered murine models with NHEJ deficiency are characterized by the onset of very aggressive pro-B cell lymphomas harboring oncogenic, V(D)J recombination-driven, Ig gene chromosomal translocations when introduced on a TP53 KO tumor accelerator background ^57^.

The occurrence of V(D)J recombination driven chromosomal translocations involving Ig and TCR genes in the context of NHEJ deficiencies highlights the imperious necessity of maintaining broken DNA ends in close proximity during DNA repair, which is also true in case of accidental DSBs introduced by genotoxic agents. The emergence of single molecule analyses coupled to cryo-EM studies in the recent years allowed a major breakthrough in our understanding of how DNA end synapsis operates to cope with DSBs by the NHEJ apparatus. These studies, which are reviewed in two very recent manuscripts ^11, 12^, mainly focused on the Ku/DNAPK-cs, XLF, PAXX, XRCC4, and DNA-Lig4 factors, established the existence of 2 major synapsis forms, Long Range (LR) and Short Range (SR), operating successively. The Ku70/80 heterodimer plays a central role in both synapses, notably through the recruitment of several DNA repair factors via their Ku binding motifs (KBMs). This is in particular the case for PAXX, MRI, and XLF. During V(D)J recombination, a third synapsis form ensures that the broken DNA coding-ends are maintained in closed proximity. This synapsis is composed of the recombinase factors RAG1 and RAG2 themselves, which remain on-site as the so-called post cleavage complex (PCC) once the DSB has been introduced at recombination signal sequences (RSS). How the PCC interacts with both LR and SR remains an interesting issue.

Our identification of a *bona fide* KBM within the c-Terminus of RAG2 suggests that, apart from its activity during initiation of the V(D)J recombination, RAG2 can be envision as participating in the DNA repair phase *per se* just like the other KBM-containing factors such as PAXX and MRI, which also exert functional redundancy with XLF. Interestingly, we showed that the functional redundancy indeed relies on the KBMs in these various partners. The C terminus region of RAG2 is required to stabilize the synapse formed by the PCC as demonstrated with a particular mutant of RAG2 (RAG2^c/c^), lacking this region. Although RAG2^c/c^ mice are not immuno-compromised, they display important genomic instability that translates into the development of thymic lymphomas when crossed onto the TP53 KO genetic background ^20^. One can argue that the PCC at least acts as a backup synapsis in case either LR or SR synapses is defective. It can be hypothesized that when XLF is absent, impacting the formation of the SR synapsis, the PCC then functionally replaces the SR synapsis, explaining the functional redundancy established between the C terminus of RAG2 and XLF. Indeed, XLF being a critical NHEJ factor both *in vivo* and *in vitro*, cells that are XLF deficient are severely sensitive to any genotoxic agent causing DSBs but in contrast to our expectation, XLF deficient lymphoid cells appeared proficient for V(D)J recombination and XLF KO mice did not present the SCID phenotype. Moreover, XLF/TP53 doubly deficient animals did not demonstrate onset of pro-B cell lymphomas, arguing for a specific backup system bypassing the loss of XLF for the repair of RAG1/2 driven prDSBs. It appeared that RAG2 itself, more specifically its C-terminal region (RAG2-CT), is key in the functional redundancy with XLF, indeed restricting this backup system to the resolution of V(D)J associated prDSBs as opposed to general genotoxic inflicted DSBs. As a consequence, RAG2^c/c^ mice, which express a RAG2 protein devoid of its C-terminus, present a T-B-SCID phenotype when crossed onto XLF KO animals, owing to abortive V(D)J recombination in lymphoid precursors^22,23^. Moreover, RAG2^c/c^/XLF animals develop B cell lymphomas in the context of TP53 deficiency. We report that RAG*2^c/c^/MRI^-/-^* and *RAG2^c/c^/PAXX^-/-^* cells show no detectable V(D)J recombination defect, arguing that RAG2 c-terminus (R2CT) is epistatic with PAXX and MRI. Since like RAG2^c/c^, PAXX and MRI are also functionally redundant with XLF, this suggests that PAXX and MRI contribute to the alternate PCC-mediated synapsis that can rescue the lack of SR synapsis in case of XLF deficiency.

We propose that specific genome guardian mechanisms have co-evolved with processes generating programmed DSBs as to mitigate deleterious consequences in case of DNA repair weakness. In this study we highlighted the central role of Ku as a master factor safeguarding the genome integrity during V(D)J recombination through the combinatorial functional redundancy of factors bearing KBMs, including RAG2. This central role of Ku can be further extended, notably during PGR in *paramecia* ^19^. Indeed, the presence of Ku even constitutes a prerequisite for PiggyMac driven initiation of PGR. In contrast to PGR, although we found that RAG2 does interacts with Ku in the absence of proficient V(D)J reaction, this interaction is not mandatory to initiate V(D)J recombination as shown with RAG2^c/c^ mice not displaying V(D)J recombination deficiency, but may ensure that other Ku interacting DNA repair factors will be available “on-site” as soon as the DNA break is introduced.

## Methods

### Cell culture

All cell-lines were grown in media consisting of DMEM (GlutaMax, GIBCO) supplemented with 10% heat-inactivated fetal bovine serum (FBS, Sigma), 100 U/mL penicillin/streptomycin (GIBCO), 1 mM sodium pyruvate (GIBCO), 2 mM L-glutamine (GIBCO), 1X nonessential amino acids (GIBCO), and 55 µM 2-Mercaptoethanol (GIBCO) at 37 °C.

### Preparation of retroviral supernatants

Retroviral supernatants were prepared by transfection of Platinum-E cells ^58^ with pMSCV- *v-abl* plasmid encoding for Abl ^38^, pMSCV-Bcl2-IRES-puro plasmid encoding for Bcl2 and puromycin selectable marker ^59^, pMX-RSS-GFP/IRES-hCD4 (pMX-INV) reporter plasmid encoding for the cell surface marker human CD4 ^21, 38^ and all the cDNA expression constructs from Empty vector, RAG2^WT^, RAG2^Δ1^, RAG2^Δ2^, XLF^WT^, XLF^ΔC^, MRI^WT^, MRI^ΔNΔC^, PAXX^WT^, and PAXX^ΔC^ using jetPRIME (Polyplus), according to manufacturer’s protocols, harvested 48 h and 72 h after transfection, snap frozen and conserved at −80°C.

### Generation of *v-abl* transformed pro-B cell-lines

Total bone marrow from 3–5-week-*PAXX*^-/-^ old mice was cultured and infected with a retrovirus encoding *v-abl* kinase to generate immortalized pro-B cell-lines ^60^. *v-abl* transformed pro-B cell-lines were then transduced with pMSCV-Bcl2-puro retrovirus ^59^ to protect them from *v-abl* kinase inhibitor-induced cell death.

### V(D)J recombination assays

The pMX-INV V(D)J recombination substrate was introduced in pro-B cell-lines through retroviral infection and cells that had integrated the recombination substrate were enriched based on hCD4 expression ^21, 38^. For V(D)J recombination assay, *v-abl* transformed Bcl2/pMX-INV infected pro-B cells (1 million per ml) were treated with 0.5 uM of the *v-abl* kinase inhibitor ABLki (PD180970, Sigma) and assayed for rearrangement by FACS analysis of GFP expression at 0, 48, 72 or 96 h. V(D)J recombination efficiency was scored as the percentage of GFP positive cells among hCD4-positive cells ((hCD4-PE antibody; 1/200 dilution). Data were acquired on a BD-LSR Fortessa (BD Biosciences) and analyzed using FlowJo software.

### CRISPR/Cas9 editing of wild type and *XLF^-/-^ v-abl* Bcl2 pro-B cells

CRISPR/Cas9-mediated gene editing was used to create a number of different *v-abl* pro-B cell-lines deficient for NHEJ factors. We utilized WT (LD12095 G1.20) *v-abl* pro-B cell-lines ^23, 26^ to generate *RAG2^-/-^, MRI^-/-^, v-abl* pro-B cell clones; WT (LD12096 A2) cell-lines were used to generate *RAG1* catalytic dead cells; *XLF^-/-^* (LD12004 C5) cell-lines were used to generate *XLF^-/-^/MRI^-/-^*, *XLF^-/-^/PAXX^-/-^*, , *XLF^-/-^/RAG2^-/-^ v-abl* pro-B cell clones. One of the *XLF^-/-^/MRI^-/-^ v-abl* pro-B cell clones were used to disrupt PAXX gene to generate *XLF^-/-^/MRI^-/-^/PAXX^-/-^ v-abl* pro-B cell clones. *RAG2^c/c^* (LD12019) were used to generate *RAG2^c/c^/MRI^-/-^*, *RAG2^c/c^/XLF^-/-^, RAG2^c/c^/PAXX^-/-^*. One of the *MRI^-/-^ v-abl* pro-B cell clones were used to disrupt PAXX gene to generate *MRI^-/-^/PAXX^-/-^ v-abl* pro-B cell clones. Briefly, Guide RNAs (gRNAs) targeting *RAG1, RAG2, MRI, PAXX and XLF* genes were synthesized as 20 bp fragments (Eurofins) with the following sequences summarized in Table S1, and cloned into pSpCas9(BB)-2A-GFP (pSpCas9(BB)-2A-GFP (PX458) was a gift from Feng Zhang (Addgene plasmid # 48138 ; http://n2t.net/addgene:48138 ; RRID:Addgene_48138). The guide sequence containing Cas9 vector and pmCherry-C1 empty vector (3 ug each) were mixed transfected in to 2 million *v-abl* cells using NEPA21 Electroporator were mixed and then introduced with a NEPA21 electroporator (Nepa Gene Co., Ltd., Chiba, Japan). After 48 h of transfection, cells were selected based on GFP status using BD FACS Aria II SORP cell sorter and colonies were allowed to form from single cells. Genotypes of all cell-lines used were authenticated by sanger sequencing. Clones for which inactivating mutations were identified on both alleles were selected for functional analysis through V(D)J recombination assays. A minimum of two independent knock-out clones were assayed for V(D)J recombination and compared to the non-edited mother *v-abl* pro-B cell-line.

### PCR analysis of Igκ V(D)J recombination products

Endogenous *Vk*_6-23_/*Jk1* CJs were amplified as previously described ^38^. A total of 500 ng of genomic DNA was amplified using pkJa2 and pk6d primers for CJ. Serial 4-fold dilutions of this reaction were amplified using pkJa2 and pk6c primers for CJ ^23^.

### Anti-GFP pull-down and mass spectrometry analysis

*v-abl* pro-B cells stably expressing either GFP-R2CT or GFP were treated with ABLki to induce RAG expression for 72 h, cells were harvested and the pellets were directly extracted for 30 min with buffer-A (20 mM Tris-HCl, pH 7.5, 300 mM NaCl, 1 mM Na_3_Vo_4_, 1 mM NaF, 1% NP40, 1 mM phenylmethylsulfonyl fluoride (PMSF), 10% glycerol and supplemented with complete EDTA-free protease inhibitor cocktail (Roche # 04693116001). After incubation added 1 volume buffer-B (20 mM Tris-HCl, pH 7.5, 1 mM Na_3_Vo_4_, 1 mM NaF, 1% NP40, 1 mM phenylmethylsulfonyl fluoride (PMSF), 10% glycerol and complete EDTA-free protease inhibitor cocktail. Following ultra-centrifugation at 14,000 rpm for 30 min at 4 °C, the supernatant was incubated with anti-GFP nanobodies coupled to magnetic beads, with end-to-end mixing for 16 h at 4 °C ^45, 48^. In some experiments, EtBr (50 mg/mL) or benzonase (25 units/ mL, Novagen) was added in the extraction buffer. Bound complexes were washed at least three times (10 min) with buffer A + B (1:1 v/v). Finally, beads with the bound complexes were directly re-suspended in SDS loading buffer (50 mM Tris-HCl (pH 6.8), 2% SDS, 300 mM 2-mercaptoethanol, 0.01% bromophenol blue, and 10% glycerol), fractionated by reducing and denaturing SDS-PAGE, and analyzed by Coomassie staining and mass spectrometry.

For LC-MS/MS Analysis, beads were washed trice with 100 μL of 25 mM NH4HCO3. Finally, beads were resuspended in 100 μL of 25 mM NH4HCO3 and digested by adding 0.2 μg of trypsine/LysC (Promega) for 1 h at 37 °C. Samples were then loaded into custom-made C18 StageTips packed by stacking one AttractSPE® disk (#SPE-Disks-Bio-C18-100.47.20 Affinisep) and 2mg beads (#186004521 SepPak C18 Cartridge Waters) into a 200 µL micropipette tip for desalting. Peptides were eluted using a ratio of 40:60 MeCN:H2O + 0.1% formic acid and vacuum concentrated to dryness. Peptides were reconstituted in 10µl of injection buffer (0.3% TFA) before nano-LC-MS/MS analysis. Online chromatography was performed using an RSLCnano system (Ultimate 3000, Thermo Scientific) coupled to an Orbitrap Fusion Tribrid mass spectrometer (Thermo Scientific). Peptides were trapped on a 2 cm nanoViper precolumn (i.d. 75 μm; C18, Acclaim PepMapTM 100, Thermo Scientific) with buffer A (2:98 MeCN:H2O in 0.1% formic acid) at a flow rate of 4.0 µl/min over 4 min. Separation was performed on a 50 cm nanoViper column (i.d. 75 µm, C18, Acclaim PepMapTM RSLC, 2 μm, 100Å, Thermo Scientific), regulated to a temperature of 55 °C with a linear gradient from 5% to 25% buffer B (100% MeCN in 0.1% formic acid) at a flow rate of 300 nl/min over 100 min. Full-scan MS was acquired using an Orbitrap Analyzer with the resolution set to 120,000, and ions from each full scan were higher-energy C-trap dissociation (HCD) fragmented and analyzed in the linear ion trap.

For protein identification, the data were searched against the Mus Musculus UP000000589_10090 database (downloaded 01/2019 containing 22266 entries) using Sequest HT through Proteome Discoverer (v2.2). Enzyme specificity was set to trypsin and a maximum of two missed cleavages sites were allowed. Oxidized methionine and N-terminal acetylation were set as variable modifications. Maximum allowed mass deviation was set to 10 ppm for monoisotopic precursor ions and 0.6 Da for MS/MS peaks. The resulting files were further processed using myProMS ^61^ v3.10. False-discovery rate (FDR) was calculated using Percolator ^62^ and was set to 1% at the peptide level for the whole study. Label-free quantification was performed using peptide extracted ion chromatograms (XICs), computed with MassChroQ ^63^ v2.2.1. For protein quantification, XICs from proteotypic peptides shared between compared conditions (TopN matching) were used. Missed cleavages and modifications were not allowed for quantification. Median and scale normalization at peptide level was applied on the total signal to correct the XICs for each biological replicate (N=3). To estimate the significance of the change in protein abundance, a linear model (adjusted on peptides and biological replicates) was performed, and p-values were adjusted using the Benjamini–Hochberg FDR procedure. Proteins with at least 2 distinct peptides in 3 replicates of a same state, a 2-fold enrichment and an adjusted p-value ≤ 0.05 were considered significantly enriched in sample comparisons. Proteins unique to a condition were also considered if they matched the peptides criteria. The mass spectrometry proteomics data have been deposited to the ProteomeXchange Consortium (http://proteomecentral.proteomexchange.org) via the PRIDE partner repository ^64^ with the dataset identifier PXD045640 (username, password: BtCe3Rnw).

### Antibodies, SDS-PAGE and immunoblotting

Whole cell extracts were obtained by washing 30 million G1-arrested wild type abl pre-B cells in phosphate-buffered saline (PBS) before lysis in SDS loading buffer (50 mM Tris-HCl pH 6.8, 2% SDS, 300 mM 2-mercaptoethanol, 0.01% bromophenol blue and 10% glycerol). SDS-PAGE was performed using the 4-20% SDS-PAGE (Bio-Rad). Immunoblotting was performed as described previously ^45^. Antibodies utilized in this study include Anti-Ku70 (ThermoFisher # MA5-13110), Anti-Ku80 (ThermoFisher # MA5-12933), Anti-RAG1, and Anti-β-Actin (sc-47778). All the antibodies were used at dilution of 1: 2,000.

### Generation of PAXX, XLF, MRI, RAG2, WT and mutants

The human RAG2 C-terminal coding sequence (aa351 to 527) was amplified by PCR using RAG2^CT^ Xho-F and RAG2^CT^ HindIII-R primers (Table. S2) from pcDNA 1.1 hRAG2-myc (JPV #229) and ligated into a pEGFP-C2 vector between XhoI and HindIII sites. The GFP-RAG2^CT^ fragment was then released by digesting with SnaBI and SmaI and cloned in to linearized pMSCV-pBabeMCS-IRES-RFP with SnaBI and AleI. Similarly, the GFP insert from pEGFP-C2 was release by digesting with AfeI and SmaI and cloned in to linearized pMSCV-pBabeMCS-IRES-RFP. The murine MRI coding sequence was amplified by PCR using primers MRI_F and MRI_R (Table. S2) from the GenEZ ORF Clone: OMu09342D and ligated into a retroviral vector pMSCV. All the mutants were generated by Q5 Site-Directed Mutagenesis Kit (see Table S2 for primer sequences). All the plasmid constructs were verified by sanger sequencing.

### Purification of Ku full-length and Ku^ΔC^ heterodimers

Expression and purification of Ku heterodimers for calorimetry was carried out as described previously ^37^. The full-length human Ku heterodimer (Ku70 1-609, Ku80 1-732) was cloned into a Multibac vector. The construct has a 10x His-tag and a TEV site on the N-terminus of Ku80 and no-tag on Ku70. Protein production was initiated in Sf21 insect cells by infection with the baculovirus stock at MOI of 5×10^-3^ and cells were collected 5-6 days after the infection (3-4 days after the proliferation arrest). Cells were sonicated and the supernatant was incubated with Benzonase (300 units for 30 min at 4 °C). The Ku heterodimer was purified on a NiNTA-Agarose affinity column (Protino, Macherey Nagel) with a 1M NaCl wash step to remove DNA excess. The eluted Ku was then bound onto an anion exchange column (Resource Q, Cytiva) and eluted with a salt gradient. The final yield was about 20 mg of purified heterodimer per litre of culture. Ku^ΔC^ constructs, heterodimer deleted of its C-terminal region (Ku70 1-544, Ku80 1-551) was cloned in the same vector and produced following the same protocol as Ku^FL^ expressed in Sf21 insect cells.

### Calorimetry

Interactions between Ku^ΔC^, Ku full-length, DNA and peptides derived from RAG2 last 39 amino acids were determined by Isothermal Titration Calorimetry (ITC) using a VP-ITC calorimeter (Malvern). Prior to measurements, all solutions were degassed under vacuum. The reaction cell of the ITC (volume 1.8 mL) was loaded with Ku heterodimers at 10 µM or 20 µM concentration. Proteins were dialyzed against buffer 20 mM Tris pH8, 150 mM NaCl, 5 mM β-mercaptoethanol. The syringe (290 µL) was filled with peptides at concentration 10-fold higher than the samples in the cell (100 µM or 200 µM). The Ku heterodimer present in the cell was titrated by automatic injections of 6-10 μL of the different ligands. All binding experiments were performed at 25 °C. Enthalpy ΔH (in kcal.mol^-1^), stoichiometry of the reaction N, and association constant Ka (in M^-1^) were obtained by nonlinear least-squares fitting of the experimental data using the single set of independent binding sites model of the Origin software provided with the instrument. Control experiments were performed with peptides injected into the buffer to evaluate the heat of the dilution. The competition between RAG2-KBM and PAXX-KBM was performed by incubating Ku^ΔC^-DNA (42 bp) at 20 µM with PAXX peptide at 200 µM in the ITC cell before titration by RAG2 peptide.

### Statistics

All data were analyzed by one-way or two ANOVA with Dunnett’s multiple comparisons test using GraphPad Prism 8 (GraphPad Software Inc.). All data are reported as the median, arithmetic or geometric mean ± SD as appropriate. Data were considered significant at a P value of less than 0.05 (*p < 0.05, **p < 0.01, ***p < 0.001, ****p < 0.0001)

### Data availability statement

The mass spectrometry proteomics data have been deposited to the ProteomeXchange Consortium (http://proteomecentral.proteomexchange.org) via the PRIDE partner repository ^64^ with the dataset identifier PXD045640 (username, password: BtCe3Rnw).

## Acknowledgments

This work was supported by INCa, LNCC, AT-Europe, and “Région Ile-de-France” and Fondation pour la Recherche Médicale grants (to D.L.). SKT was supported by grants from INCa and AT-Europe. PC and PF were supported by ANR-20-CE11-0026. PC is a scientist from INSERM. JBC thanks grants ANR-20-CE11-0026, ANR-21-CE12-0019, ANR 23-CE11-0033, ANR-23-CE18-Nejin and FRISBI national infrastructure ANR-10-INBS-0005.

## Author contributions

SKT, AG, FI, VR, FD, and DL performed experiments. PF, PC, IC, and JBC contributed to conceptualization, visualization, and review and editing of the manuscript. SKT and JPV conceived the study and wrote the manuscript.

