## Supplementary material for "Ku-binding motifs in RAG2, XLF, PAXX and MRI support functional redundancy during V(D)J recombination": Supp figures: Tadi et al_suppl material_bis.pdf

### Supplemental Figures

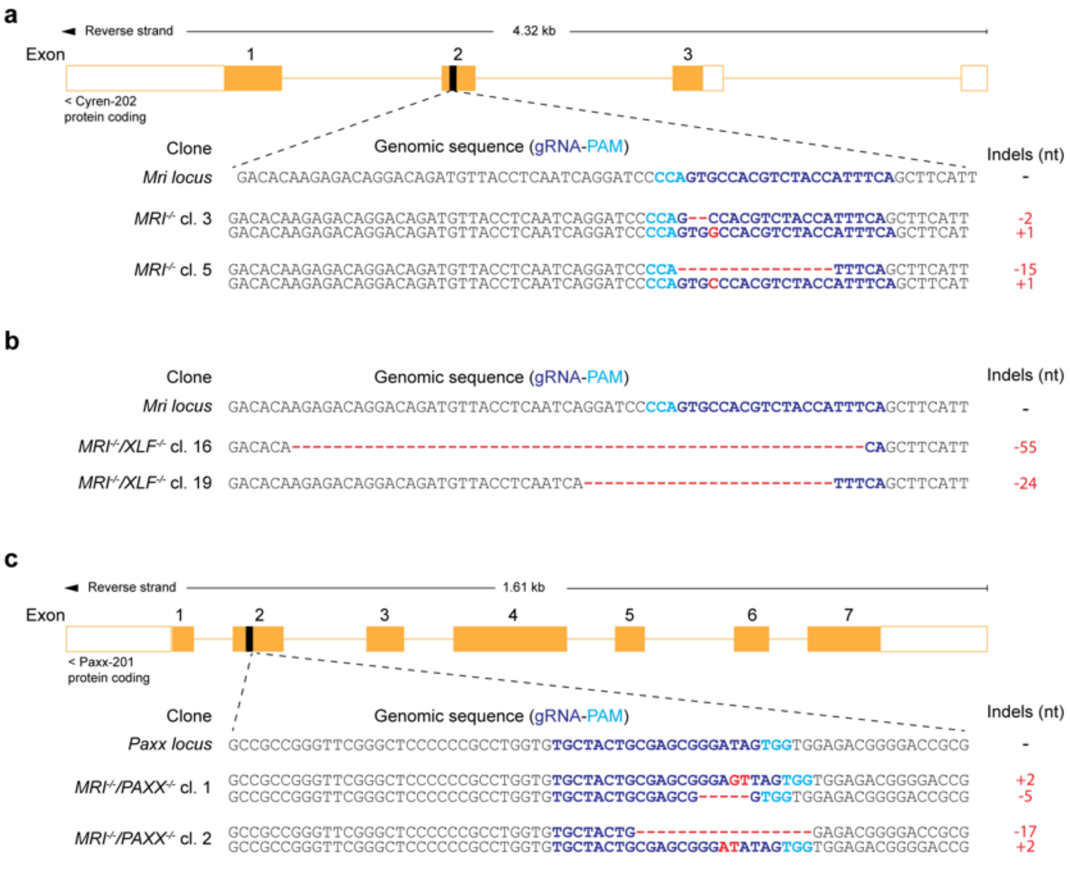


**Fig. S1**

**Fig. S1. Generation of *MRI*$^{-/-}$, *MRI*$^{-/-}$/ *XLF*$^{-/-}$ and *MRI*$^{-/-}$/ *PAXX*$^{-/-}$ *v-abl* pro-B cells using CRISPR-Cas9**. **a, b** Schematic representation of gRNA-E targeting exon 2 of *Mri/Cyren* from WT and *XLF*$^{-/-}$ *v-abl* pro-B cells to generate *MRI*$^{-/-}$, *MRI*$^{-/-}$/ *XLF*$^{-/-}$ *v-abl* pro-B cells respectively. **c** Schematic representation of gRNA#2 targeting exon 2 of *Paxx* were used to disrupt the gene function to generate *PAXX*$^{-/-}$ *v-abl* pro-B cells. Representing 2 clones of each used in the study and loss of gene functions were verified by Sanger sequencing analysis.

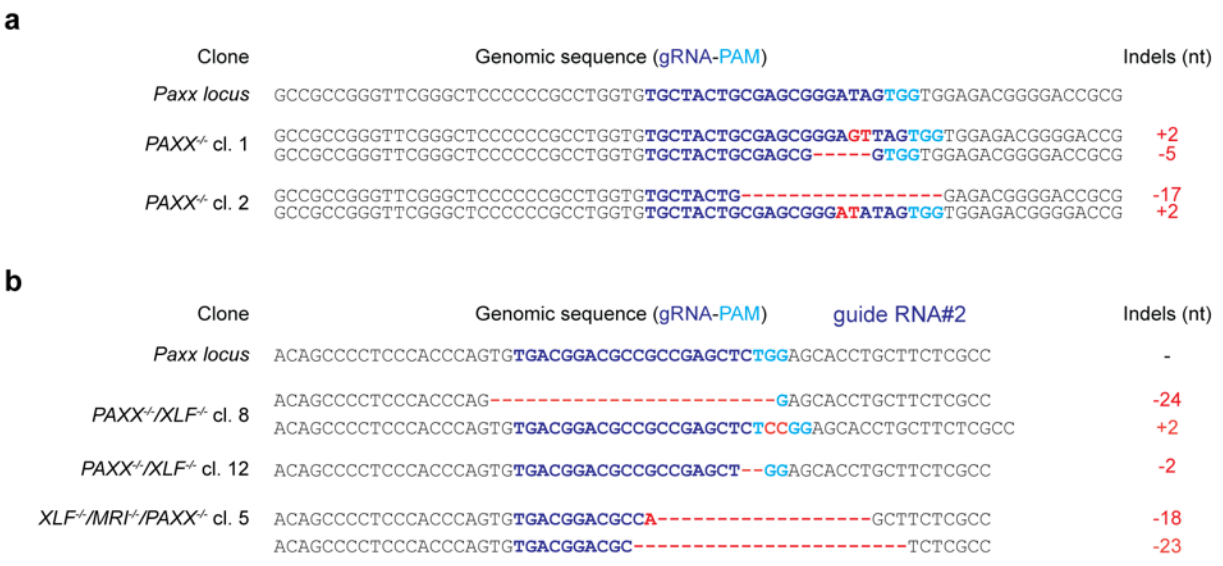


**Fig. S2**

**Fig. S2. Generation of *PAXX*$^{-/-}$, *PAXX*$^{-/-}$/ *XLF*$^{-/-}$ and *XLF*$^{-/-}$/*MRI*$^{-/-}$/ *PAXX*$^{-/-}$ *v-abl* pro-B cells using CRISPR-Cas9**. **a, b** Schematic representation of gRNA and gRNA#2 targeting exon 2 of *PAXX* from WT, *XLF*$^{-/-}$ and *MRI*$^{-/-}$/ *XLF*$^{-/-}$ *v-abl* pro-B cells to generate *PAXX*$^{-/-}$, *PAXX*$^{-/-}$/ *XLF*$^{-/-}$ and *MRI*$^{-/-}$/ *XLF*$^{-/-}$/ *PAXX*$^{-/-}$ *v-abl* pro-B cells respectively. Sanger sequencing analysis was used to select the best knockout clones. Representing 2 clones of each from *PAXX*$^{-/-}$ and *PAXX*$^{-/-}$/ *XLF*$^{-/-}$ and a single clone from *MRI*$^{-/-}$/ *XLF*$^{-/-}$/ *PAXX*$^{-/-}$ *v-abl* pro-B cells used in the study.

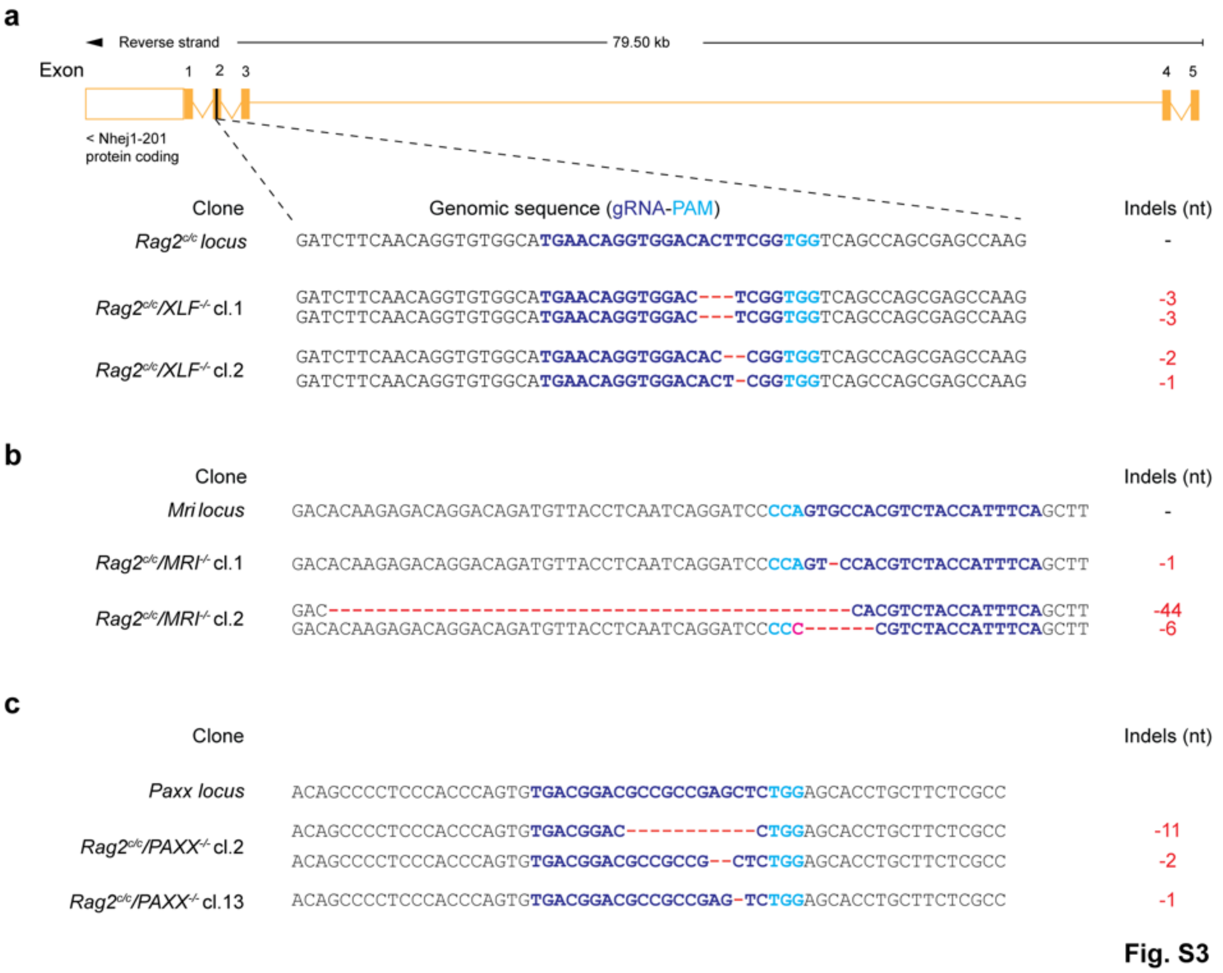


**Fig. S3. Generation of *Rag2$^{c/c}$/ XLF$^{-/-}$*, *Rag2$^{c/c}$/ MRI$^{-/-}$*, *Rag2$^{c/c}$/ PAXX$^{-/-}$ v-abl* pro-B cells using CRISPR-Cas9**. Schematic representation of gRNA#2 targeting exon 2 of *XLF*, gRNA-E targeting exon 2 of *MRI/CYREN* and gRNA#2 targeting exon 2 of *PAXX* were used to disrupt the gene function. Sanger sequencing analysis was used to select the best knockout clones. Representing 2 clones of each used in the study.

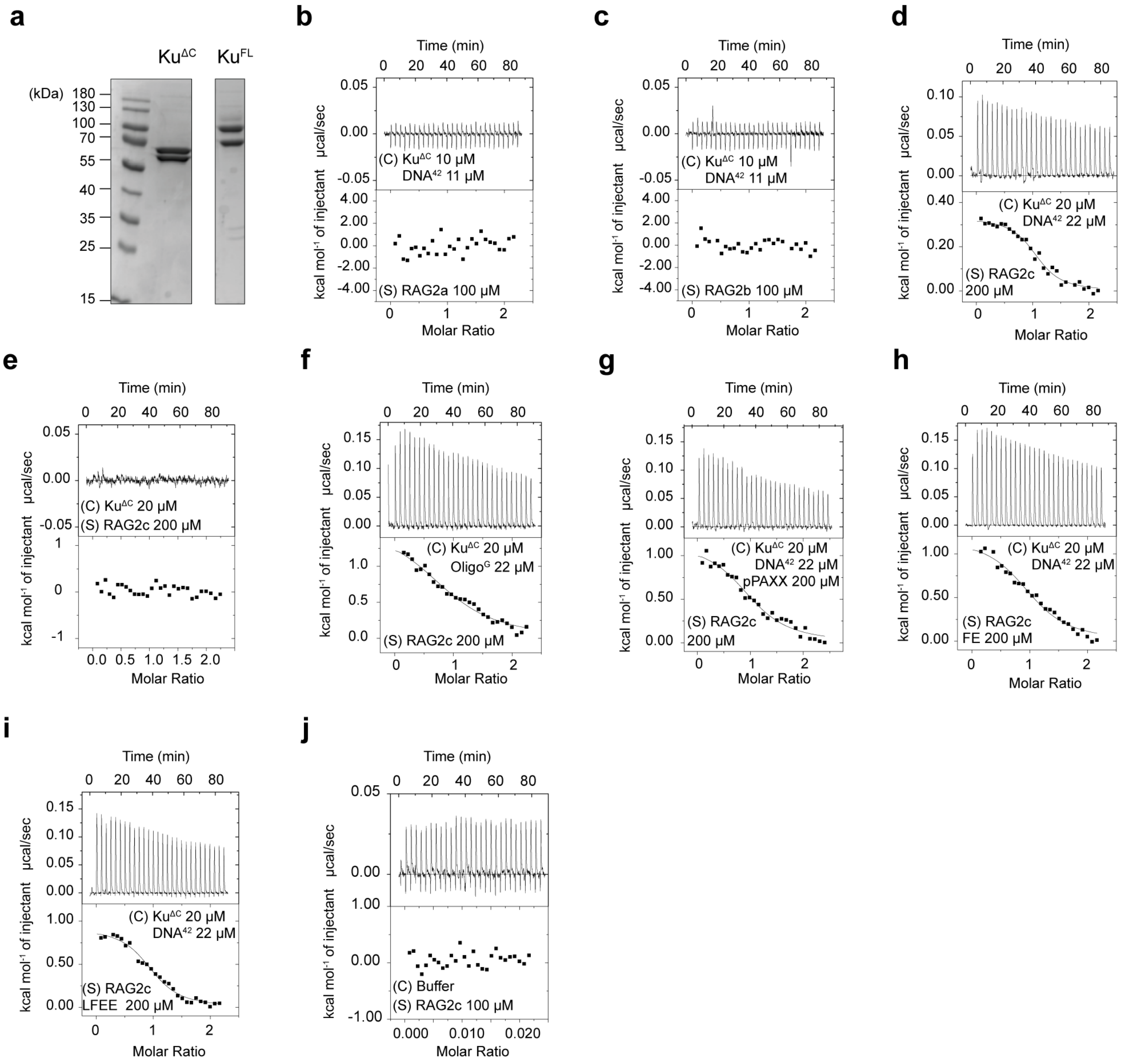


**Fig. S4**

**Fig. S4. Representative ITC thermograms, Related to Table 1**

See accompanying Table 1 and experimental procedures in the main manuscript for conditions. C = cell, S = syringe.

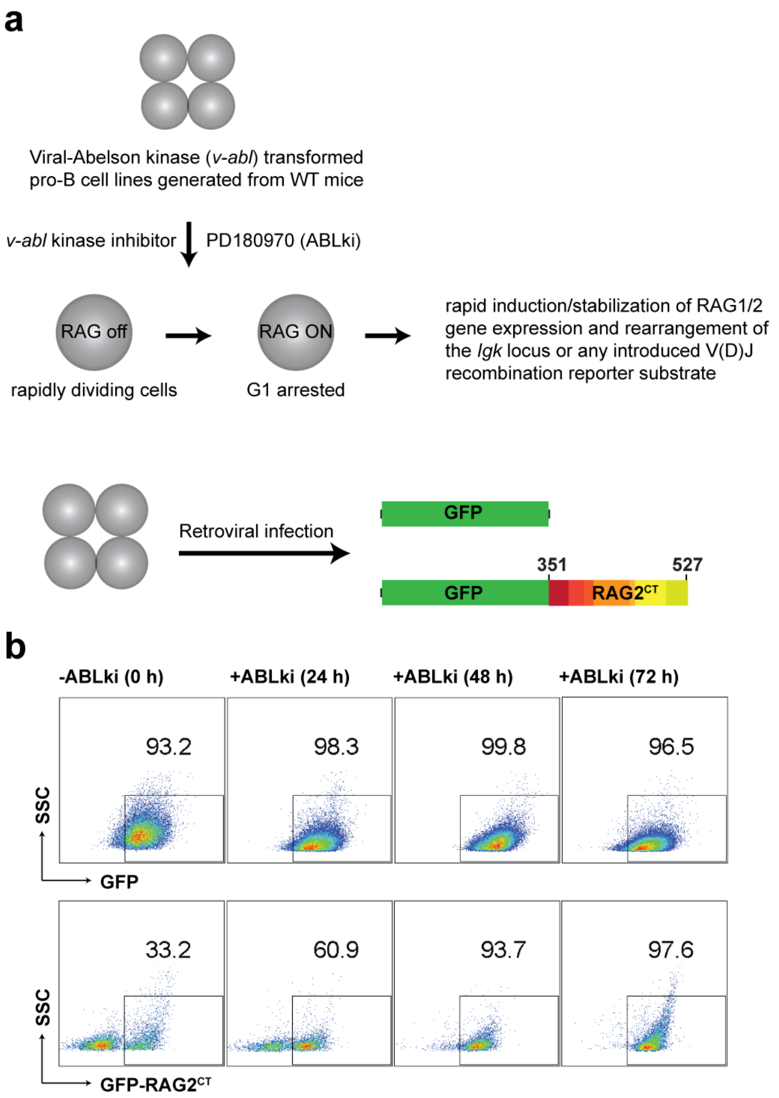


**Fig. S5**

**Fig. S5. Experimental strategy to perform RAG2$^{CT}$ interactome analysis in *v*-bl pro-B cells. a** Stimulation of *v-abl*-transformed pro-B cell lines with 0.5 µM ABLki (PD180970, Sigma) induces RAG expression, arrests cells in the $G_1$ phase and rearrangement of endogenous *Igk* locus. *v-abl*-transformed pro-B cell lines stably expressing GFP, GFP-RAG2$^{CT}$ by retroviral infection used in this study. **b** Flow cytometric analyses of GFP and GFP-RAG2$^{CT}$ expression in *v-abl* pro-B cells treated with ABLki for the indicated lengths of time.

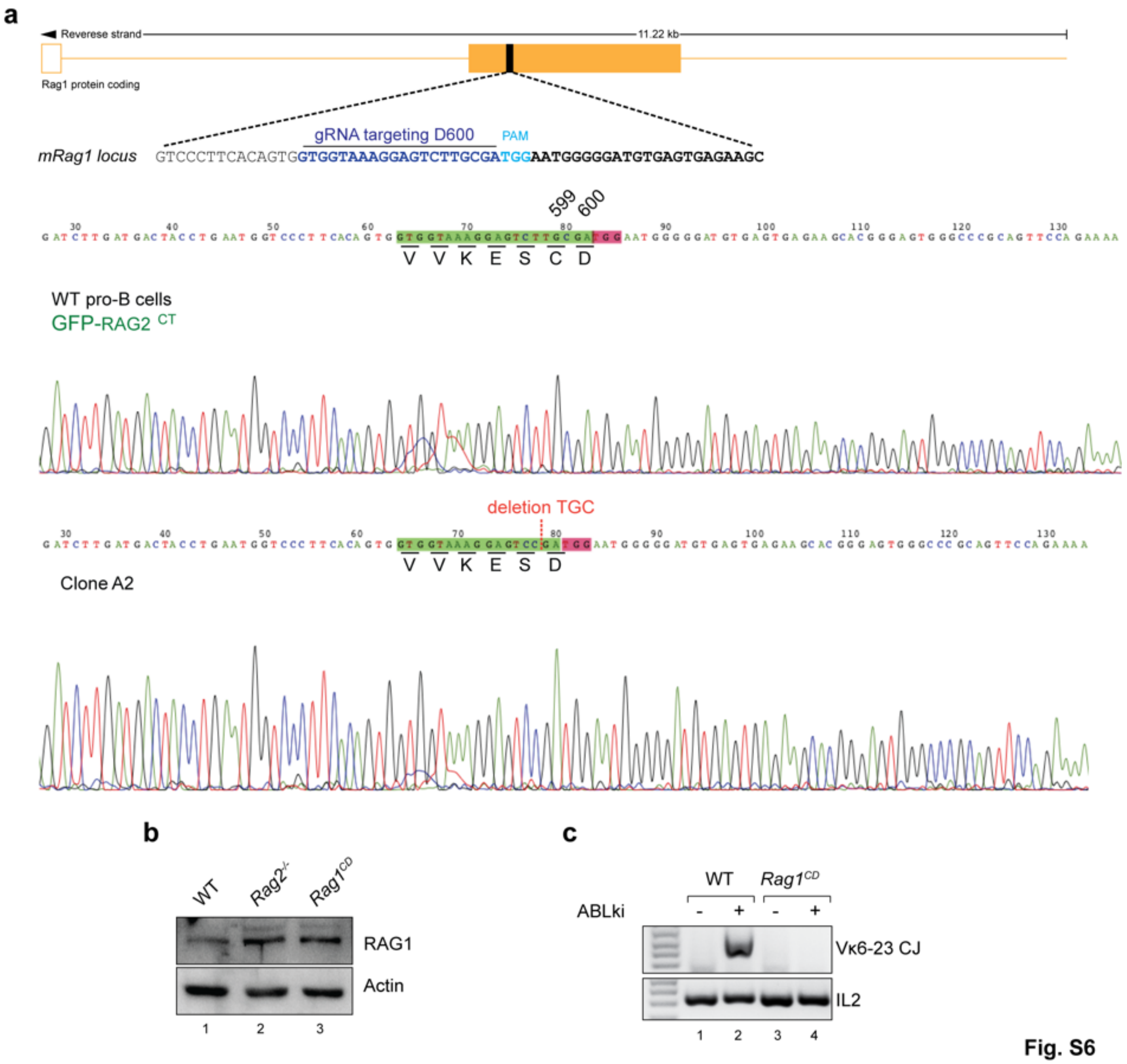


Fig. S6

**Fig. S6. GFP-RAG2$^{CT}$ interactome in *v*-bl pro-B cells. a** Schematic of the generation *Rag1$^{CD}$ v-abl* pro-B cells using CRISPR-cas9 with a gRNA targeting D600 reside. Sanger sequencing showing loss of C599 residue in *Rag1$^{CD}$ v-abl* pro-B cells. **b** Western blotting for RAG1, RAG2 and *β*-actin in lysates from WT, *Rag2$^{-/-}$, Rag1$^{CD}$ v-abl* pro-B cells. **c** PCR analysis of IgkV$_{6\text{–}23}$-J$_1$ CJ from indicated *v-abl* abl pro-B cell lines treated for 72 h with ABLki. *Il2* gene PCR was used as a loading control.

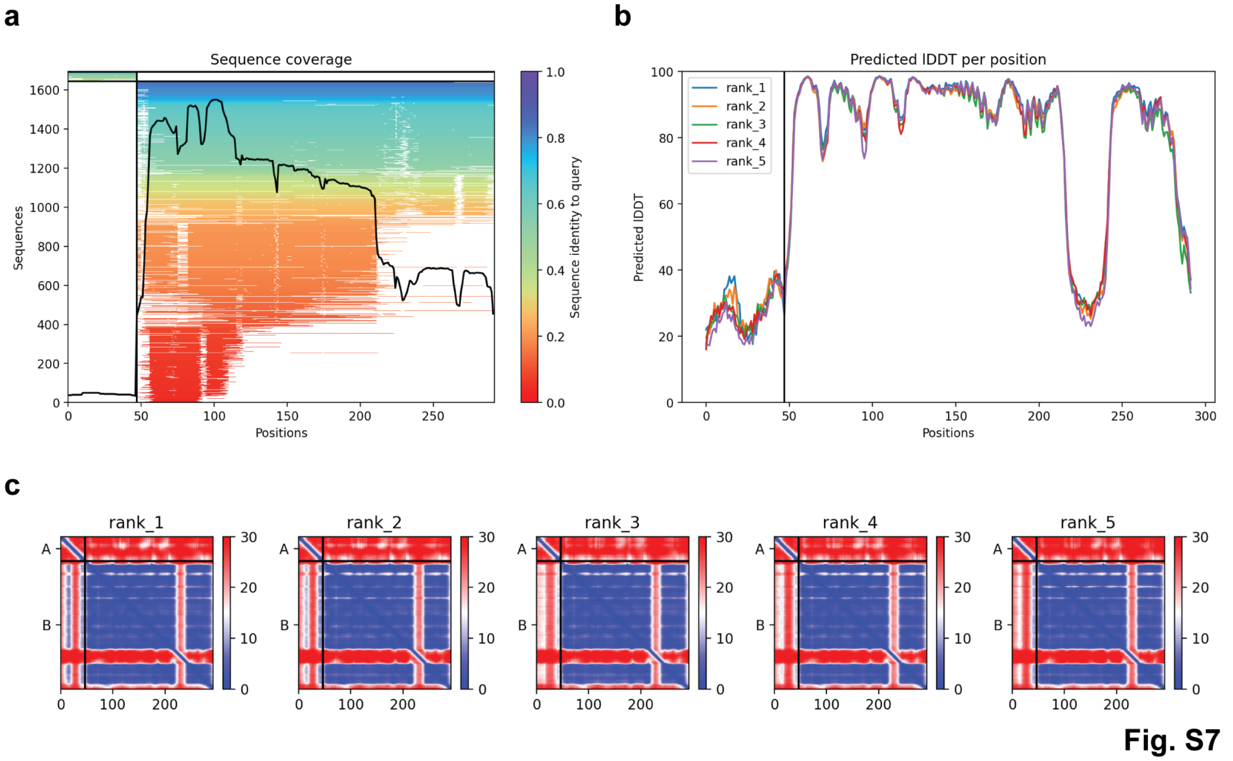


**Fig. S7. AlphaFold model of R2CT/Ku interaction**

Details of the AF2 multimer modeling: predicted iDDT per position, sequence coverage and PA. The RAG2 peptide is followed by the ku80 vWA domain.

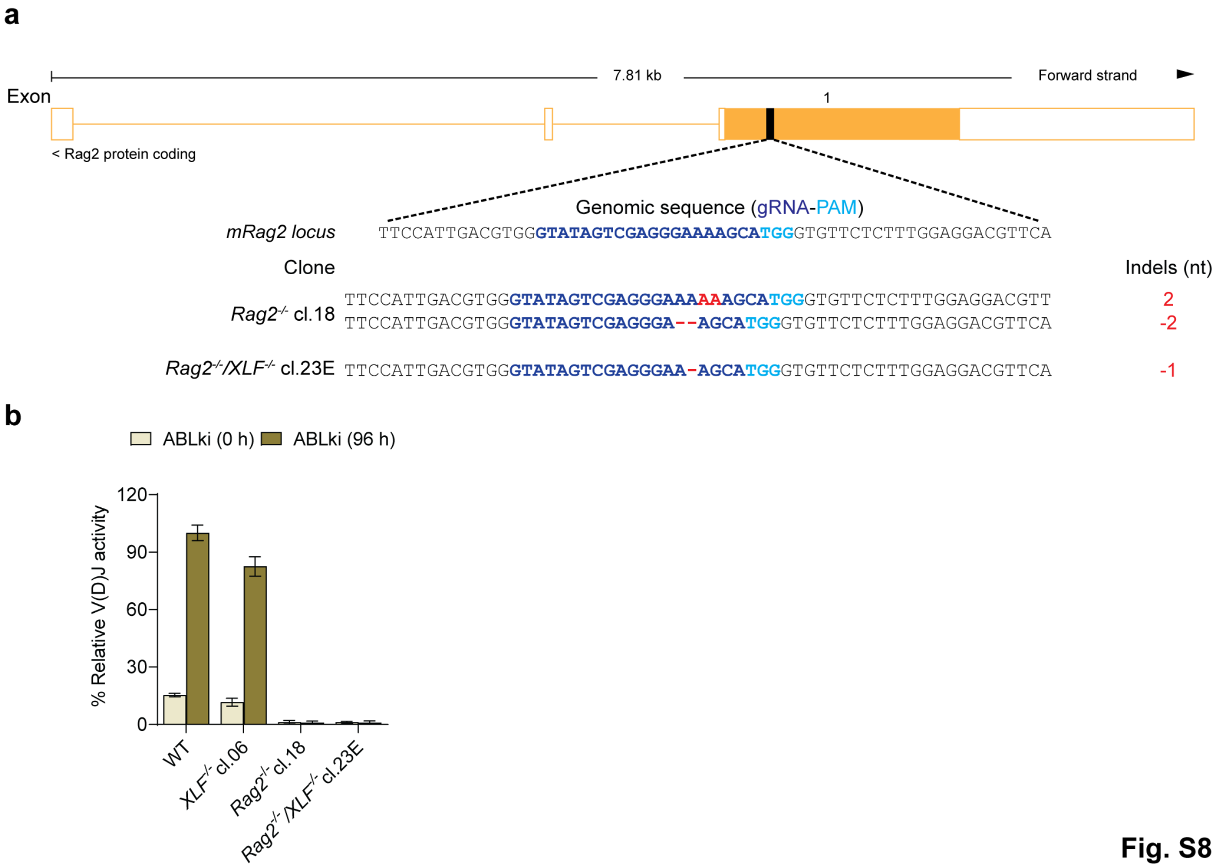


**Fig. S8**

**Fig. S8. Generation of *Rag2*$^{-/-}$ and *Rag2*$^{-/-}$/ *XLF*$^{-/-}$ *v-abl* pro-B cells. a** Schematic for the disruption of *Rag2* gene in *v-abl* pro-B cells using CRISPR-cas9 to generate *Rag2*$^{-/-}$ and *Rag2*$^{-/-}$/*XLF*$^{-/-}$. Sanger sequencing showing disruption of both the alleles in *Rag2*$^{-/-}$ and homozygous deletion of one nucleotide in *Rag2*$^{-/-}$/*XLF*$^{-/-}$. **b** Flow cytometric analyses of percent V(D)J activity in WT, *XLF*$^{-/-}$, *Rag2*$^{-/-}$ and *Rag2*$^{-/-}$/*XLF*$^{-/-}$ *v-abl* pro-B cells treated with ABLki for 0 and 96 h. Histograms represent means (± SD) of at least three independent experiments.

**Table S1. List of oligonucleotides used to construct CRISPR-Cas9 gRNAs.**

| N° | Name of the oligonucleotide | Sequence (5'-3') |
|---|---|---|
| 1 | mRag2 (gRNA-B)-Up | GTATAGTCGAGGGAAAAGCA |
| 2 | mRag2 (gRNA-B)-Low | TGCTTTTCCCTCGACTATAC |
| 3 | mRag1 (gRNA-A)-Up | GTGGTAAAGGAGTCTTGCGA |
| 4 | mRag1 (gRNA-A)-Low | TCGCAAGACTCCTTTACCAC |
| 5 | mCyren (gRNA-E)-Up | GTGAAATGGTAGACGTGGCAC |
| 6 | mCyren (gRNA-E)-Low | GTGCCACGTCTACCATTTCAC |
| 7 | mPaxx (gRNA#1)-up | TGCTACTGCGAGCGGGATAG |
| 8 | mPaxx (gRNA#1)-down | CTATCCCGCTCGCAGTAGCA |
| 9 | mPaxx (gRNA#1)-up | TGACGGACGCCGCCGAGCTC |
| 10 | mPaxx (gRNA#2)-down | GAGCTCGGCGGCGTCCGTCA |
| 11 | mXlf (gRNA#2)-up | TGAACAGGTGGACACTTCGG |
| 12 | mXlf (gRNA#2)-down | CCGAAGTGTCCACCTGTTCA |

**Table S2. List of oligonucleotides used for ITC, Q5 site directed mutagenesis and to PCR amplify V(D)J recombination products.**

| N° | Name of the oligonucleotide | Sequence (5'-3') |
|---|---|---|
| 1 | DNA-42_up | GAACGAAAACATCGGGTACGAGGACGAAGACTGACCACGACA |
| 2 | DNA-42_down | TGTCGTGGTCAGTCTTCGTCCTCGTACCCGATGTTTTCGTTC |
| 3 | DNA 30/15_up | GATCCCTCTAGATAT |
| 4 | DNA 30/15_down | CGGATCGAGGGCCCGATATCTAGAGGGATC |
| 5 | $RAG2^{CT}$ XhoI_F | CTCGAGCCGCTGAAGATGATACTAATGAAGAGC |
| 6 | $RAG2^{CT}$–HindIII_R | AAGCTTCTAATCAAACAACCTTCTAAGAAAGGATT |
| 7 | MRI_F | GACGTGGAAGAAAACCCCGGTCCTGAAATGGAAACCCTAAAATCCAAG |
| 8 | MRI_R | CAGTGTGATGGATATCTGCAGTTAGAACTAACTGAAAAATATTTCCCTGACGTACTTCAACGCGT |
| 9 | R2 Delta1_F | CCCTCCGTAAAAAAGGTTCTGGAAAAAATCTTGACT |
| 10 | R2 Delta1_R | GTCGACTGCAGAATTCGAAGCTTCTA |
| 11 | MRI DeltaN_F | ATGACAGCCCCTGTGGAT |
| 12 | MRI DeltaN_R | TTCAGGACCGGGGTTTTC |
| 13 | MRI DeltaC_F | TAGTTCTAACTGCAGATATCCATC |
| 14 | MRI DeltaC_R | CTCCTTCTCCTCCTCCGG |
| 15 | R2 Delta2_F | TAGGTCGACCTGCAGGCA |
| 16 | R2 Delta2_R | AGTGTGTAGAGCTCTTGCTATC |
| 17 | XLF Delta_F | TAAGCGGCCGCGACTCTA |
| 18 | XLF Delta_R | CTGAGGTCTCTGCAGAGGG |
| 19 | pkJa2 | GCCACAGACATAGACAACGGAA |

| | | |
|---|---|---|
| 20 | pk6d | GAAATACATCAGACCAGCATGG |
| 21 | pk6c | GTTGCTGTGGTTGTCTGGTG |
